# Migration load, competition, and metabolic trade-offs shape spatial divergence through eco-evolutionary dynamics

**DOI:** 10.1101/2025.11.04.686607

**Authors:** David M. Ekkers, Sergio Tusso, Marina C. Rillo, Oscar P. Kuipers, G. Sander van Doorn

## Abstract

Evolutionary models predict that trade-offs related to resource utilization promote local specialization and diversification in spatially heterogeneous environments. However, diversification is also influenced by ecological feedback mechanisms - such as frequency-dependent resource use - and their interaction with migration. To investigate how spatial variation affects diversification, we conducted an evolution experiment exposing populations of *Lactococcus cremoris* to divergent selection on two carbon sources: fructose and galactose. Using a continuous culture system, we partitioned these resources into two isolated patches (allopatry), two patches with constant migration between them (parapatry), and one fully-mixed patch (sympatry). Resource specialization and subsequent phenotypic diversification evolved in all treatments, revealing a metabolic trade-off between fructose and galactose utilization. Compared to treatments without migration, spatial variation produced a distinctive temporal pattern: migration initially delayed specialization on galactose but later accelerated it. This temporal pattern emerged from eco-evolutionary feedbacks operating through two mechanisms: (i) early in the experiment, higher population density in the better-suited fructose patch increased the influx of maladapted immigrants into the galactose patch, slowing local adaptation; however, (ii) as strains adapted locally to galactose, population density increased in the galactose patch and density asymmetries weakened. Simultaneously, the competitive ability of immigrants declined as residents became increasingly specialized in their local resources, progressively reducing the effective migration rate and allowing accelerated divergence. Our experiment demonstrates how the likelihood and tempo of spatial divergence depend on the evolvability of phenotypic trade-offs and their dynamic interaction with migration load and population density.

## 1 INTRODUCTION

Evolutionary theory predicts that spatial variation in heterogeneous environments can promote adaptive diversification, resulting in the establishment of multiple genetically determined specialists, each adapted to local conditions [Amarasekare and Nisbet, 2001, Amarasekare, 2003, Jeltsch et al., 2013]. Two key requirements for the emergence of phenotypic diversification are predicted by theoretical modeling: (i) the presence of fitness trade-offs that prevent the evolution of a generalist strategy with high performance across all conditions; and (ii) local density regulation generating frequency-dependent selection to maintain a global polymorphism of locally adapted, diverged phenotypes [Levins, 1968, Neuman, 1970, Van Tienderen, 1991, Wilson and Yoshimura, 1994, Kisdi, 2001, Egas et al., 2004]. Whereas the first requirement highlights the role of constraints inherent to the internal organization of the organism, the second underscores the importance of eco-evolutionary feedback mechanisms acting within the local environment.

The ecological context of adaptive diversification in response to spatial variation has received considerable attention, particularly in the theoretical literature, because the potential for diversification may depend on seemingly subtle differences in the mode of operation of ecological mechanisms. For instance, it is difficult to say in general how migration between local subpopulations will influence the likelihood or rate of diversification, as the effect of migration may depend on factors such as population density, standing genetic variation or asymmetries in dispersal [Gavrilets, 2000, Ravigné et al., 2009, Tusso et al., 2021]. Gene flow can accelerate specialization by increasing genetic variation in local populations, providing new material for selection to act on [Ching et al., 2013], or by setting the stage for competition between locals and immigrants that can increase the rate of local adaptation [Osmond and de Mazancourt, 2013]. At the same time, local populations can be swamped by an inflow of locally maladapted genes carried by incoming immigrants, which may potentially delay local specialization by interfering with the fixation of beneficial mutations [Bolnick and Nosil, 2007, Alzate et al., 2017]. The effective rate of migration is expected to depend on the progression of diversification as well, as it is determined not only by the rate of immigration, but also by the extent to which immigrants can contribute to future generations within the new habitat. In the presence of resource utilization trade-offs, increased adaptation to one local habitat automatically implies a reduced performance of immigrants in the other, especially when the residents there have also adapted to their local conditions [Johansson, 2008, Siepielski et al., 2016, Alzate et al., 2017].

The ecological mechanism of local density regulation can strongly influence the likelihood of adaptive diversification. Conditions for the maintenance of ecological polymorphism are most favorable when local adaptation is mediated by soft selection [Wallace et al., 1968, Levene, 1953, Bell et al., 2021]. Under soft selection, local population density is regulated by relative fitness within each local patch, allowing subpopulations to persist even when containing locally maladapted phenotypes. In contrast, hard selection operates on absolute fitness values, independent of local conditions. This scenario may often be more realistic: in many cases, local population density is positively correlated with the degree of local adaptation. Consequently, unequal resource use arising from pre-adaptation or constraints may translate into an (initial) asymmetry in habitat suitability. By causing unequal population sizes between habitats, such an asymmetry would result in source-sink dynamics, in which relative migration load is decreased in the source population and increased in the sink population [Bisschop et al., 2019]. If sufficiently strong, this effect is theoretically expected to undermine local adaptation to the initially unsuitable habitat; evolution would then only lead to increased specialization to a single habitat (local resource specialization) rather than to the emergence of a polymorphism of resource specialists [Ravigné et al., 2009]. These theoretical predictions require empirical validation under controlled conditions.

Systematic empirical tests of spatial diversification theory remain relatively scarce, partly due to the technical difficulties of identifying the constraints that give rise to phenotypic trade-offs and tracking how they are navigated by evolving populations [Hashemi et al., 2023, Amarasekare, 2003]. Many resource utilization trade-offs emerge as system-level properties in the context of complex metabolic pathways and regulatory networks [Schuetz et al., 2012, Hashemi et al., 2023, Polz and Cordero, 2016, Navid et al., 2019], making it hard to pinpoint exactly their underlying molecular mechanisms [Winterbach et al., 2013]. To overcome this challenge, we investigated the dynamics of divergent adaptation in a microbial evolution experiment, using the lactic-acid bacterium *Lactococcus cremoris* MG1363 (formerly *L. lactis* subsp. *cremori*) as our model system, and by exerting selection on its carbon utilization pathways. *L. cremoris* has a relatively simple carbon metabolism, which has been characterized in detail owing to its ubiquitous use in fermented dairy products, both in traditionally and industrially produced food products [Neves et al., 2005, Kok et al., 2017]. The molecular basis of resource utilization trade-offs between fructose and galactose in this model system has been characterized in detail [Ekkers et al., 2020], making it an exceptional model system to study the molecular mechanisms underlying phenotypic trade-offs and diversification in spatially heterogeneous environments.

Like in many other bacteria, the carbon metabolism of *L. cremoris* consists of a central glycolytic backbone (for the conversion of intermediate metabolites) that is shared between most carbon sources (Fig. 1a). As a result, whenever the optimization of the metabolism for a single carbon source requires the tuning of a flux rate in this core glycolytic pathway, it can have side effects (positive or negative) on the rate of growth on alternative carbon sources. Negative effects constrain efficient co-utilization of multiple carbon sources and reveal resource utilization trade-offs. These negative effects are expected when the optimization of the pathway for one sugar requires a change in the net flux direction of a reversible reaction relative to another sugar. Such incompatibilities may arise, for example, when the pathways for alternative carbon sources feed into the core glycolytic backbone at different entry points, either upstream or downstream of important anabolic offshoots. We selected fructose and galactose as alternative carbon sources because these sugars have conflicting net resource fluxes, making their optimization mutually constrained (Fig. 1a). These two sugars create, therefore, a resource utilization trade-off that is mechanistically well characterized.

**FIGURE 1.**
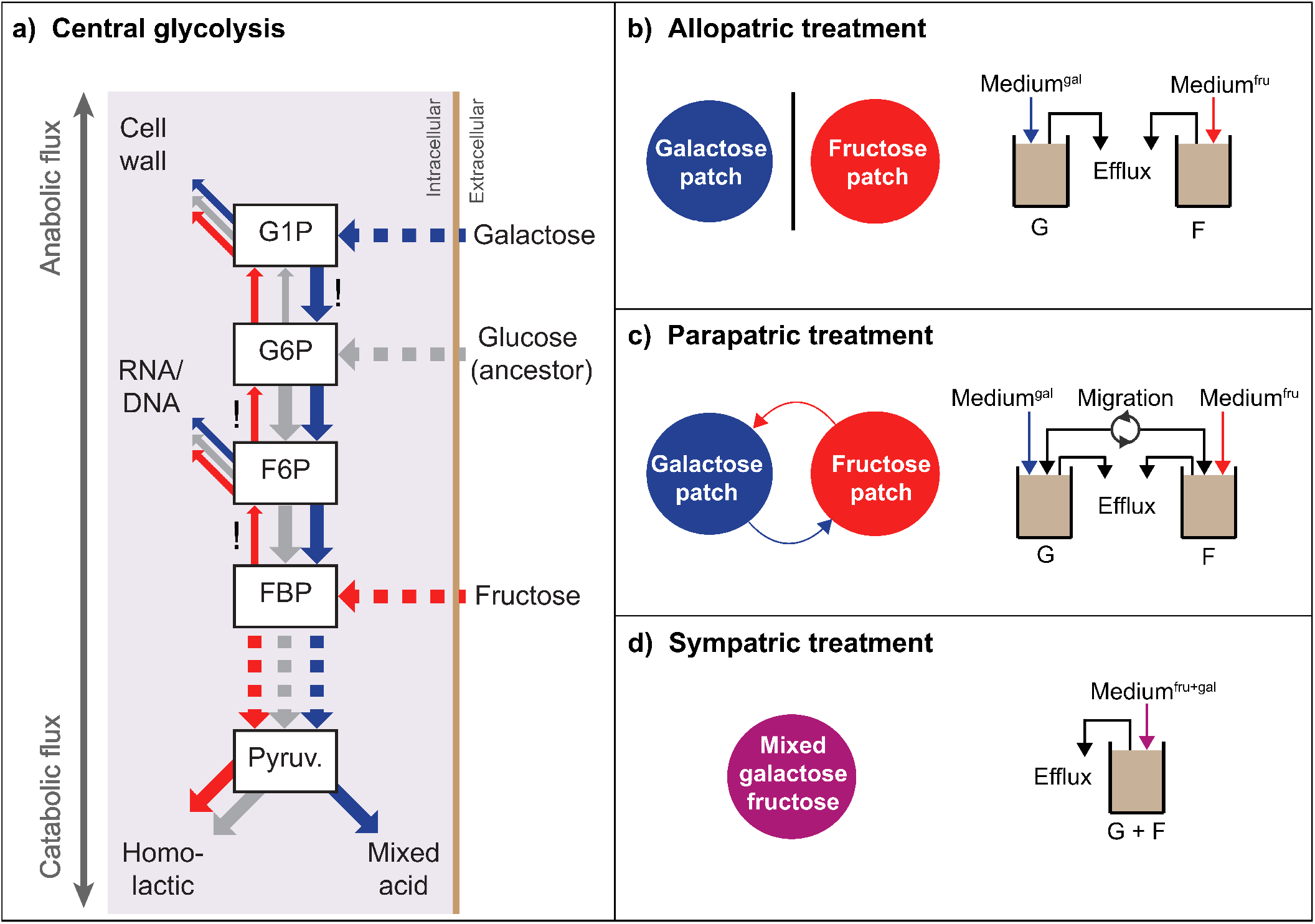
**(a)** Schematic representation of the metabolic architecture of the central carbon metabolism of *L. cremoris*. Metabolites are shown as squares: glucose 1-phosphate (G1P), glucose 6-phosphate (G6P), fructose-6-phosphate (F6P), fructose-biphosphate (FBP), pyruvate (Pyruv.). Arrows show the net direction of metabolic flux for glucose (gray), fructose (red), and galactose (blue); arrow width indicates the relative flux rate. Exclamation marks denote flux reversals compared to the ancestral glucose-adapted metabolism: upstream FBP *→* F6P and F6P *→* G6P for fructose and downstream G1P *→* G6P for galactose. For a complete glycolytic flux map, see [Ekkers et al., 2022]. Other panels illustrate the experimental evolution regimes: **(b)** allopatry, with two isolated patches: one supplemented with galactose (G) medium (Medium^*gal*^) and the other with fructose (F) medium (Medium^*fru*^); **(c)** parapatry, with galactose and fructose patches connected by migration (with a flux of 5% relative to the dilution rate of the chemostat), and **(d)** sympatry, with a fully mixed resources patch (Medium^*fru*+*gal*^).

Besides the challenges related to the molecular basis of trade-offs, the interpretation of experimental studies of adaptive diversification across heterogeneous environments is further complicated by the fact that fitness differences between alternative resource-utilization strategies are ecologically intertwined with density regulation. This association implies that fitness trade-offs are dynamic and responsive to changes in resource utilization and availability [Brown et al., 2004, Burger et al., 2019]. To confront this challenge, we exposed the experimental *L. cremoris* populations to three selection regimes that differed with respect to the ecological mechanism that regulate resource availability: *(1)* in one treatment, populations were grown in isolated resource patches, established by separate cultures supplemented with either fructose or galactose, with no migration between them (allopatric treatment; Fig. 1b); *(2)* in another, we created resource heterogeneity in space by coupling the two resource patches (fructose or galactose) by means of exchange of migrants between cultures (parapatric treatment; Fig. 1c), and *(3)* in a third treatment, cells were grown in a well-mixed culture with continuous availability of both resources (sympatric treatment; Fig. 1d). Both the selection pressure for higher growth rate and the migration between resource patches were controlled by a density-controlled continuous culture system [Ekkers et al., 2020], which allowed for a steady-state culturing regime. Such a regime avoids the large temporal fluctuations of selective pressure that are inherent to serial transfer protocols [Gresham and Dunham, 2014], allowing us to specifically test the effects of spatial variation in the absence of temporal variation. After the evolution experiment, we *(i)* characterized the phenotypes of the evolved strains in each treatment, *(ii)* compared the evolved strains with each other and with the ancestral strain, *(iii)* performed genetic, transcriptomic and enzymatic analyses to link the phenotypic adaptations with newly emerged genetic variants, and lastly *(iv)* compared density differences and competitive ability between migrants and resident populations to assess source-sink dynamics. To our knowledge, this is the first study to integrate an eco-evolutionary analysis of spatial diversification with a mechanistic understanding of resource utilization trade-offs and their response to selection at the molecular level.

## 2 MATERIAL AND METHODS

### 2.1 Experimental procedures

Prolonged evolution experiments employing continuous cultures are susceptible to biofilm formation. We therefore chose *L. cremoris* MG1363 with its low biofilm-forming properties as opposed to more readily used bacterial model systems such as *Escherichia coli, Pseudomonas fluorescens* or *Bacillus subtilis*. The evolution experiment was performed using a custom-built continuous-culture system [Ekkers et al., 2020]. *L. cremoris* was grown in 60 ml of chemically defined medium for prolonged cultivation (CDMPC) [Price et al., 2019]. Cultures were grown anaerobically (25 ml/min N2 headspace flow) under continuous dilution and stirring (330 rpm) at 30°C and maintained at a constant pH of 6.5 by dynamically pumping NaOH solution (30% wt/v) into the bioreactors. Spatial variation in the treatment with two connected resource patches (parapatric treatment) was created by using two bioreactors, one only supplemented with fructose medium and the other one with only galactose medium (Fig. 1c). Migration between the two patches was induced by exchanging a fixed volume of culture between the two bioreactors with a peristaltic pump (*Reglo Digital*). The migration rate was set to 5% of the chemostat dilution rate (m = 0.05 D, where D is the dilution rate in h^*−*1^). ensuring that migration per generation remained constant throughout the experiment. Two control treatments were run in parallel: allopatric with isolated galactose and fructose patches (i.e., without migration; Fig. 1b) and sympatric treatment with both resources together (Fig. 1d). We used media supplemented with either galactose (1% wt/v), fructose (0.5% wt/v) for the allopatric treatment, and for the sympatric treatment, a mixture of galactose (0.5% wt/v) and fructose (0.25% wt/v) as sole carbon source(s). For each of the treatments, we ran four replicates in parallel.

The ancestral, glucose-adapted *L. cremoris* exhibited much slower growth on galactose than on fructose. The unequal amounts of fructose and galactose (1:2 wt/v ratio) that were provided in the culture media compensated partially for the observed initial asymmetry in growth rate. To further balance the population sizes across treatments at the start of the evolution experiment, the initial dilution rate was set at the maximum dilution rate of the ancestor population for each resource environment, resulting in a starting dilution rate of 0.2 for the parapatric and allopatric-galactose treatments, and at 0.3 for the allopatric-fructose and sympatric treatments. Throughout the experiment, samples were drawn daily from each bioreactor to measure OD_600_ and preserve population-level (chemostat-level) glycerol stocks. To keep selection pressure constant despite increased growth rates, the dilution rates of the bioreactors of each treatment (i.e., allopatric-fructose, allopatric-galactose, parapatric, and sympatric) were continuously adjusted daily to maintain population density at steady state between 0.5 < OD_600_ < 1.0. Given that the dilution rate approximates the per-capita growth rate of the culture at steady state, we monitored its change throughout the experiment for signs of evolutionary adaptation and selected three timepoints (T1, T2, and T3) for subsequent phenotypic analysis. The experiment was run for 38-58 days, depending on the treatment. The number of generations per treatment was calculated from the daily dilution rate as (*D × t*)/ ln(2), where *D* is the dilution rate and *t* is time in hours, corresponding to an estimated 759-1603 generations of evolution (Supp. Fig. S1).

### 2.2 Library construction of evolved strains

A library of single-genotype strains was constructed by plating population samples on a fructose and a galactose plate, and picking six colonies per plate. In total, 12 strains were sampled from each replicate population from each treatment for each timepoint, yielding a single-strain library of 432 single strains. To build this library, we diluted population samples from the glycerol stock 500 times and plated 50 *μ*l of the diluted cell suspension on F-CDMPC (fructose 1%) agar plates for allopatric-fructose and parapatric-fructose patches, on G-CMDPC (galactose 1%) plates for the allopatric-galactose and parapatric-galactose patches, and on both types of agar plates for the sympatric treatment. We picked six colonies from each plate. Each colony was then grown separately overnight at 30°C without shaking in 2 ml of F-CDMPS or G-CDMPC (identical to the agar plate it was derived from), and glycerol stocks were prepared from these cultures.

### 2.3 Growth curve analysis

We measured growth rates on fructose and galactose for all 432 strains from the library after pre-culturing them overnight in F-CDMPC and G-CDMPC. We diluted the -80°C stocks 100 times in PBS and used 1 *μ*l of this cell suspension to inoculate 100 *μ*l of fresh fructose or galactose chemically defined medium for prolonged cultivation (CDMCP) adjusted to pH 6.5. Each strain was grown with three replicates in 384-well plates (Greiner Bio-one 781906) under anaerobic conditions (VIEWseal Greiner Bio-one) at 30°C in a plate reader (Tecan F200). Estimates of the instantaneous growth rate were obtained by performing local linear regression analyses on the growth curve data, using a sliding window of five data points (measurements were taken every 10 min). After eliminating noisy data from the initial growth phase (OD < 0.16), we determined the maximum growth rate from the regression curves. We then averaged the three replicate values of the maximum growth rates for each strain on each sugar. It is important to note that for the sympatric populations, growth measured on each sugar separately should not be interpreted as a direct estimate of absolute fitness in the mixed-resource environment. Rather, these assays decompose adaptation into fructose-specific and galactose-specific performance components, allowing divergence along the focal trade-off axis to be compared across treatments.

### 2.4 Phenotypic clustering analysis

We used a model-based clustering method to identify and classify the different phenotypic groups that evolved during the evolution experiment. The clustering was performed on the maximum growth rates of the single strains on each of the two sugars. The performance of each strain was plotted as a two-dimensional coordinate: F^*max*^ (maximum growth rate on fructose) on the y-axis and G^*max*^ (maximum growth rate on galactose) on the x-axis. The growth rate coordinates in the F^*max*^-G^*max*^ space were clustered per timepoint per treatment. To estimate the number of clusters (phenotypic groups), we applied normal (Gaussian) mixture models and a maximum likelihood approach using the R package *mclust* (version 5.4, Scrucca et al. 2016). To consistently apply the same clustering model to all the treatments and timepoints, we selected the ‘EVV’ model, which allows clusters of ellipsoidal shape (i.e., bivariate Gaussian distributions) with different covariance structures (i.e., different orientations). This model was either the best model selected by the Bayesian Information Criterion (BIC, maximum likelihood corrected for model complexity) or yielded the same number of clusters as the best model selected by BIC independently for each timepoint and treatment. We limited the maximum number of clusters to three because we were only interested in the main phenotypic groups in the F^*max*^-G^*max*^ space (i.e., a fructose specialist, a galactose specialist, and a generalist). For a complete visualization of the clusters see Supp. Fig. S2.

### 2.5 Metabolic profile table

A set of three representative phenotypes was selected from each phenotypic cluster of each treatment (including 6 genotypes from the spatial, 6 from the mix, and 6 from the isolated control treatments). Growth measurements on fructose, glucose, mannose, and galactose (1% wt/v) were conducted in the same way as had been done previously for galactose and fructose (see description under phenotypic clustering above). Besides measuring growth curves, we also determined yield by quantifying the maximum optical density of each culture.

### 2.6 DNA sequencing and analysis

#### 2.6.1 Strain selection

We sequenced 72 monoculture strains by sampling two strains from each phenotypic cluster in two out of four replicates for each timepoint for each treatment. Besides monocultures, we also sequenced 60 populations consisting of the population samples extracted from each bioreactor at each timepoint.

#### 2.6.2 DNA isolation, sequencing, and bio-informatic analysis

The isolation of genomic DNA of the population-level glycerol stocks, as well as from the single-strain library was performed as described by Johansen and Kibenich [1992]. Full genome resequencing was performed using IlluminaHiSeq (GATC) with a mean coverage of 300x, and read length and insert size of 150 bases. To characterize genetic variation across the strains, adaptors were removed with *cutadapt* 1.18 [Martin, 2011], and pair reads were filtered and trimmed based on quality scores using *trimmomatic* 0.36 [Bolger et al., 2014] and *FastQC* 0.11.5. Filtered reads were then mapped to the reference genome (*Lactococcus lactis* subsp *cremoris* v. MG1363) using BWA 0.7.13 [Li and Durbin, 2009]. Duplicated reads were removed, and local realignment was performed using *picard* 2.18.5, increasing the maximum number of reads to 100,000 per base. Coverage values per base were calculated using *SAMtools* 1.9 [Li et al., 2009]. Across the sequenced samples, the average whole-genome read depth was 557*×*, with a minimum of 162*×* and a maximum of 825*×* (Supp. Fig. S6). Genotype calling was done with *FreeBayes* 0.9.10 [Garrison and Marth, 2012], setting a minimum mapping quality of 20 and ploidy level of 1. This procedure results in a list of genetic variants that includes single nucleotide polymorphisms (SNPs) and small insertions and deletions (INDELs). To consider only the genetic variants that appeared in the course of the experiment, the genetic variants of each sample were contrasted with the variants observed in the ancestral strain (timepoint T0). Only the genetic variants that were observed in the evolved strains but not present in the ancestral strain were retained for subsequent analysis.

Due to variation in coverage along the genome relative to the ancestral strain, alignment files (bam files) were used to identify genomic regions with copy number variation (CNV). CNV was inferred using *CNVnator* 0.3.3 [Abyzov et al., 2011], with a value of 100 for the parameter bin size and the option ‘unique’ in order to have the correct output of quality field. In this latter case, no variants were detected in the ancestral strain. To identify the potential phenotypic effect of the resulting list of genetic variants (SNPs, INDELs, and CNV), they were annotated relative to the reference genome using *SnpEff* 4.3 [Cingolani et al., 2012]. Annotations for downstream, upstream, and interacting genes related to the CNV were not included. Based on genetic analysis, we discovered candidate variants related to galactose specialists (GS) or fructose specialists (FS), the majority of which only occurred in one strain or replicate, or occurred in both GS and FS phenotypes. For practical purposes, we set a threshold to identify the genes with the most abundantly occurring phenotype-specific variants. To do so, we only considered annotated loci that exclusively occurred in one phenotypic group (FS or GS) and with four or more variants in the same replicate or that occurred in multiple replicates. For a full list of variants and selected strains (including their variants) for functional testing, see Supp. Table S1 and S3.

To calculate the number of variants for genes that were under selection, we summed all unique variants from both the monoculture and population analysis. The frequency of the adaptive variants was calculated from read abundance based on the population sequencing data. To formally test whether repeated gene targets across independent populations exceeded chance expectations, we analyzed de novo variants from the population-level sequencing data at the lineage level. Population SNP and INDEL variants were called jointly with the ancestral strain and were retained only when they showed strong support in evolved populations while showing no appreciable support in the ancestor. Variants were then annotated relative to the reference genome and collapsed by independent lineage identity. We first tested for excess genome-wide gene-level recurrence using a permutation procedure that preserved the number of mutated genes per lineage and the mutational target size of each gene. Because the focal specialist-associated targets were not expected to recur equally across all treatments, we additionally tested whether predefined FS-associated (*fbp, pfk*) and GS-associated (*pgmA, ptnABCD, ldh*) targets were enriched in ecologically relevant subsets of lineages using one-sided hypergeometric tests. For these restricted tests, fructose-specialist targets were evaluated in the allopatric-fructose and parapatric-fructose lineages, and galactose-specialist targets in the allopatric-galactose and parapatric-galactose lineages; expanded analyses including sympatric lineages were used as sensitivity analyses. P-values were corrected using the Benjamini-Hochberg correction.

The 3D protein structure prediction from the PFK and LDH proteins was performed using *Phyre2* [Kelley et al., 2015]. *EzMol* was used to visualize the proteins, color functional domains, and mutated regions [Reynolds et al., 2018]. *Protter* was used to visualize the intermembrane mutations in the mannose PTS [Omasits et al., 2014].

### 2.7 Strain selection for functional analysis of evolved variants

A set of representative strains that contained mutations in the identified adaptive regions (*fbp, pfk, pgmA, ldh*, and *ptnAB*) was selected for further functional tests to assess the effect of the mutations. At least one strain from each phenotypic cluster of each treatment (including 4 strains from the spatial, 2 from the mix, and 4 from the isolated control treatments) was included (Supp. Table S1).

### 2.8 RT-qPCR experiments

To measure expression levels of the mutated genes compared to the ancestor, we performed RT-qPCR on the transcriptome of actively metabolizing cells. We grew the selected strains in batch (60 ml) in the bioreactor system at pH 6.5, 30°C in F-CDMPC or G-CDMPC. We harvested culture samples mid-exponentially (OD_600_ *≈* 0.45) and froze the cell pellet immediately in liquid nitrogen. We performed RNA isolation and cDNA preparation in duplo. Expression levels of each sample were measured with five replicates and normalized using expression levels of the housekeeping gene *glyA* as a reference. For primer sequences for each tested gene see Supp. Table S2.

### 2.9 Lactic acid assay

To measure the dehydrogenase activity in the GS strains we measured the relative amount of lactic acid in its metabolic waste products. Cultures were grown on CDMPC supplemented by 0,5% galactose and harvested mid-exponentially (OD_600_ 0.45). Essays were performed on supernatant with a *Megazyme L-lactic acid assay kit* and performed as stated by the manufacturer’s protocol.

### 2.10 Pfk assay

To quantify the enzymatic activity of the mutated PFK protein, we measured its ability to convert ATP to ADP. Protein extracts were prepared from mid-exponentially harvested cells grown on F-CDMPC. Essays were performed with a *MyBioSource Phosphofructokinase microplate Assay kit (catalog number MBS8243182)* and executed as stated by the manufacturer’s protocol.

## 3 RESULTS

### 3.1 Phenotypic divergence reveals two specialist strategies across treatments

At the end of the experiment, evolved phenotypes exhibited a negative relationship between growth on fructose and galactose (Fig. 2a). Two groups occupy opposite ends of this trade-off: one consisting of strains with high maximum growth rate on fructose and low on galactose (fructose specialist, FS), and one with high growth rate on galactose and low on fructose (galactose specialist, GS). These two specialist phenotypes evolved independently across all treatments, matching the respective fructose or galactose resource patch. In allopatry (hereafter A), the phenotypes form a single cluster: FS_*A*_ or GS_*A*_ according to their sugar environment (Fig. 2a). In parapatry and sympatry (hereafter P and S, respectively), a polymorphism of FS and GS evolved: FS_*P*_ in the parapatric fructose patch and GS_*P*_ in the parapatric galactose patch; and FS_*S*_ or GS_*S*_ in sympatry. Despite selection for simultaneous resource use in these two treatments, no generalist phenotype evolved (Fig. 2a), consistent with the earlier observation of a strong trade-off between the maximal growth on fructose and galactose [Ekkers et al., 2022].

**FIGURE 2.**
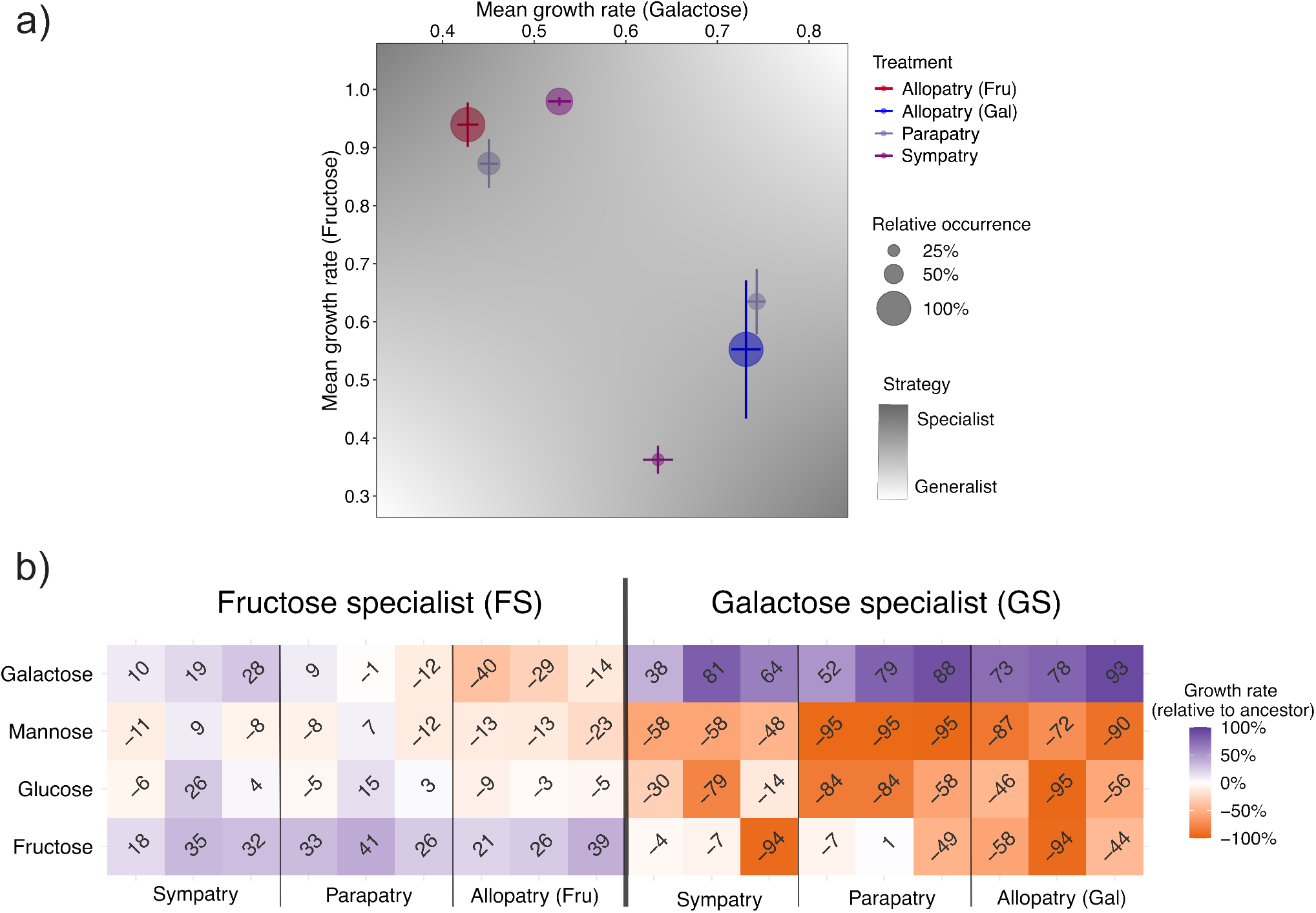
Phenotypic characterization of evolved strains. **(a)** Average maximum growth rate of the evolved populations at timepoint T3 for all treatments. Growth rates were measured in monocultures grown in CDMPC supplemented with either galactose (x-axis) or fructose (y-axis). For each treatment, 24 strains were analyzed and subsequently clustered to identify phenotypic groups. Circle size indicates the frequency of each phenotypic cluster, and error bars indicate within-cluster variance. Background shading indicates regions in phenotype space corresponding to specialist (grey) or generalist (white) strategies. Based on these clusters, we distinguish two specialist phenotypes: a fructose specialist (FS, high growth rate on fructose, low on galactose) and a galactose specialist (GS, the opposite from FS). **(b)** Comparison of the metabolic profiles of selected FS and GS strains between the sympatric, parapatric and allopatric treatments. Strains were grown in triplicate in batch culture on CDMPC supplemented with 1% (wt/v) fructose, glucose, mannose or galactose. Colors and numbers indicate relative percentual growth rate changes compared to the ancestral strain (purple = positive, orange = negative).

Metabolic analysis of specialist strains revealed additional phenotypic changes in growth rate and yield across multiple monosaccharide carbon sources compared to the ancestral strain (Fig. 2b). FS and GS showed consistent profiles of growth performance across different sugars. GS strains exhibited growth-performance trade-offs not only on fructose but also on glucose and mannose, particularly pronounced in allopatry (GS_*A*_) and parapatry (GS_*P*_). In contrast, FS strains generally lacked such trade-offs: only FS_*A*_ showed moderate performance decrease on most resources, likely due to neutral mutation rather than selection, since FS_*P*_ and FS_*S*_ did not display similar changes. Overall, FS and GS exhibited asymmetric improvements relative to the ancestor: GS increased galactose growth at the cost of fructose, glucose, and mannose growth, whereas FS did not show consistent changes. This asymmetry reflects the ancestor’s preadaptation to glucose, which enabled relatively efficient use of fructose (ancestor growth rates: glucose 1.01 h^*−1*^, fructose 0.83 h^*−1*^, galactose 0.43 h^*−1*^). Consequently, GS strains showed the largest relative improvements in growth rate (Fig. 2, Fig. 3).

**FIGURE 3.**
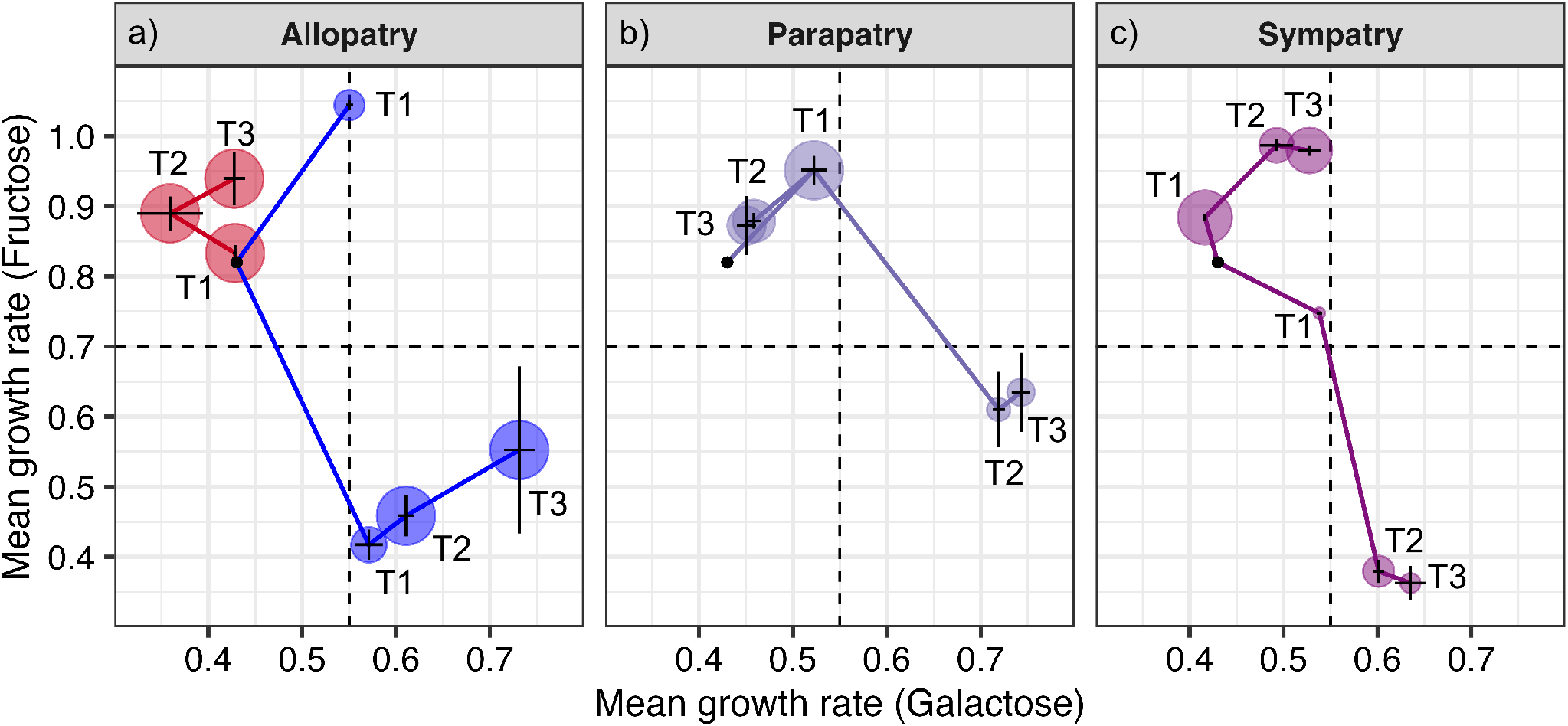
Evolutionary trajectories of the populations of the evolved strains from the three treatments: **(a)** allopatry (fructose in red and galactose in blue), **(b)** parapatry, and **(c)** sympatry. The black dot, from which the trajectories originate, indicates the phenotype of the ancestral strain. Growth rates (h^*−*1^) were calculated from evolved monocultures grown in CDMPC supplemented with either fructose (y-axis) or galactose (x-axis). The sizes of the circles indicate the relative abundance of each group of phenotypes (i.e., clusters) sampled at timepoints T1-T3, and error bars indicate variance. The dotted lines separate regions associated with specialists (upper left corner and lower right corner) and generalists (upper right corner and lower left corner).

### 3.2 Recurrent genetic targets are associated with specialist phenotypes

Whole-genome sequencing of selected strains identified five annotated genes exclusively associated with either FS (*fbp* and *pfk*) or GS (*ptnABCD, pgmA* and *ldh*) (Fig. 2c; Supp. Table S3). These genes act at key junctions of the carbon metabolism of *L. cremoris*, where flux directions were predicted to change relative to preadapted glucose metabolism (Fig. 1a). Other less frequent or unannotated variants also occurred, but were excluded from further analysis (see Methods). No coding target was exclusive to a single treatment, but recurrence of the focal mutations was not evenly distributed across all populations. Across the pooled population sequencing data, most de novo mutations remained rare: 196 of 258 unique sites were detected in only one lineage, whereas only 62 recurred across multiple lineages, and carrier observations were generally at low to intermediate allele frequency (Supp. Fig. S7).

To formally test whether repeated genetic targets exceeded chance expectations, we analyzed the population-level sequencing data using lineage-collapsed recurrence tests. Across the full set of annotated genes, recurrent hits occurred more often than expected under a null model that accounts for both lineage-specific mutational load and gene target size in both the T3 endpoint data and the full time-series dataset (Supp. Table S4). Because the focal specialist-associated mutations were not expected to recur equally across all populations, we additionally tested whether they were enriched in ecologically relevant subsets of lineages. This restricted analysis showed a clear enrichment of *pfk* mutations in fructose-specialist lineages in both the T3 and all-timepoint datasets (P = 0.0036, q = 0.036; Supp. Table S5). Signals for *fbp* and *ldh* were weaker but directionally consistent in the all-timepoint analysis (both P = 0.049), whereas *pgmA* and *ptnABCD* showed the expected bias toward galactose-specialist lineages but remained too sparse for strong statistical support.

### 3.3 Distinct metabolic changes underlie fructose and galactose specialization

A subset of the fructose specialists (FS) carried mutations in *pfk* and *fbp* (Fig. 4a), which are crucial to balance anabolic/catabolic carbon flux on fructose. The gene *fbp* encodes fructose-1,6-bisphosphatase (Fbp), which catalyzes the reverse reaction from FBP to F6P. Mutations in its upstream region increased expression two-to seven-fold compared to the ancestor (Fig. 4b). This result aligns with earlier work showing that fructose metabolism is constrained by limited anabolic flux between FBP and F6P, due to fructose’s relatively low entry point into glycolysis compared to glucose (Fig. 1a; [Looijesteijn et al., 1999]). Surprisingly, one of the FS_*S*_ strains without an *fbp* mutation (strain FS_*S*1_) showed an average 2.35-fold increase of *fbp* expression (Fig. 4b). The gene *pfk* encodes phosphofructokinase (Pfk), which catalyzes the glycolytic conversion of F6P to FBP (Fig. 4a; [Papagianni et al., 2007]), and was structurally mutated in the allosteric activation and inhibition sites (Supp. Fig. S3; Supp. Table S3; Ekkers et al. 2022). Nine out of ten independently evolved *pfk* variants involved missense mutations near allosteric regulation sites of Pfk (Supp. Fig. S3; Supp. Table S1). Expression analysis showed no change in the *pfk*-mutants (FS_*A*1_ and FS_*P*1_) and a small reduction in expression of two out of the three FS strains without a *pfk* mutation (FS_*A*2_ 20% and FS_*P*2_ 12%; Fig. 4c). Enzyme assays confirmed that Pfk catalytic activity was reduced by 25% (Fig. 4d). Interestingly, the FS strains from the parapatric and allopatric treatments that lacked *pfk* mutations still showed a small reduction in expression of *pfk* of 13% and 21%, respectively. Galactose specialists (GS) were associated with mutations in *pgmA, ptnABCD*, and *ldh*. The galactose pathway connects to the core glycolytic backbone upstream of the entry point of glucose (Fig. 1a). *pgmA* encodes phosphoglucomutase, the main bottleneck of galactose metabolism, and its upregulation promotes the catabolic flux from G1P to G6P (Figs. 1a, 4e; Neves et al. 2006, 2010). Mutations in the gene *pgmA* included variants in the upstream region of the gene and duplications of sections of the genome that included *pgmA*. The *pgmA* mutant strains displayed a two to three-fold expression increase compared to the ancestor (Fig. 4f). A strain (GS_*P*1_) without a *pgmA* mutation showed no expression change (Fig. 4f). The gene *ptnABCD* codes for the mannose phosphotransferase system (Man-PTS), which is the primary import system for glucose and mannose (Fig. 4e; Neves et al. 2005). The mutations in *ptnABCD* largely consisted of variants located in the intergenic region and the transmembrane components of *ptnC* and *ptnD* (Supp. Fig. S5). Given that six out of seven non-intergenic variants caused either a frameshift (4x), gain of stop codon (1x), or a deletion (1x) (Supp. Fig. S5, Supp. Table S3), we expect the mutations to have a deleterious effect. Expression experiments confirmed that the intergenic upstream mutations resulted in a nearly complete repression of expression (91-99%) (Fig. 4g). Lastly, *ldh* gene (lactate dehydrogenase), encodes the key metabolic enzyme that converts the reversible pyruvate-lactate reaction (Fig. 4e), the terminal step in homolactic fermentation on glucose or fructose. Five different *ldh* variants were found (Supp. Table S3; Supp Fig. S4) and, based on the quantification of metabolic waste products (lactic acid) of *ldh* mutants, we conclude that lactate dehydrogenase activity was strongly reduced in the mutated strain (GS_*A*1_, Fig. 4h).

**FIGURE 4.**
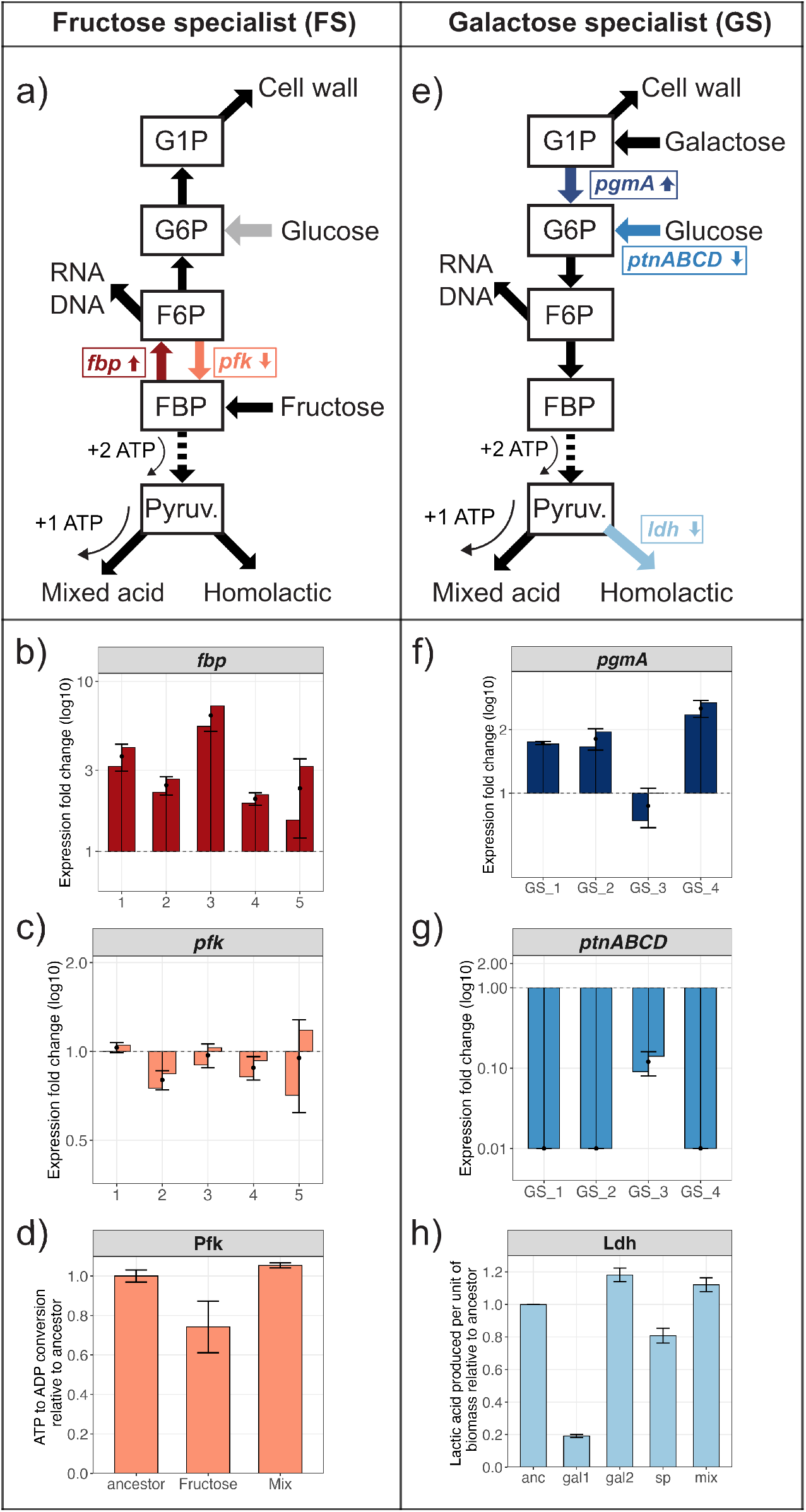
Functional validation of the genetic variants associated with the fructose specialist strains (FS) and galactose specialist strains (GS). **(a)** Mutated genes of FS and **(e)** GS mapped to the architecture of the central glycolysis. Small vertical arrows indicate measured change in either expression or enzymatic activity (up: increase; down: decrease). **(b)** Expression changes of *fbp* gene. **(c)** Expression changes of the *pfk* mutants. **(d)** Pfk activity essay. **(f)** Expression changes of the *pgmA* mutants. **(g)** Expression changes of the *ptnABCD* mutants. **(h)** Ldh activity essay. Dots indicate the average, and error bars standard deviation between the two measurements. The variants and strains that are tested are listed in the Supp. Table S1. For protein changes, see Supp. Table S3 and Supp. Figs. S3, S4, S5.

In summary, all mutations reside in locations in the metabolism where flux changes occur between the selected resource (fructose or galactose) and the preadapted resource (glucose). The functional analysis of the two evolved specialist phenotypes (FS and GS, Fig. 2) and their associated mutations and transcriptional patterns (Fig. 4) point towards two consistent functional adaptations: for FS, the metabolic flux between FBP and F6P is altered by increasing metabolic activity in the anabolic direction (*fbp*) and decreasing it in the catabolic direction (*pfk*) (Fig. 4a). The GS increased catabolic flux between G1P and G6P (*pgmA*), silenced the activity of the primary glucose sugar import system (*ptnABCD*), and decreased homolactic fermentation (*ldh*) (Fig. 4e).

### 3.4 Evolutionary adaptation is delayed in the parapatric treatment

We tracked evolutionary trajectories in each treatment by measuring growth performance on fructose and galactose at three timepoints (T1, T2, T3) spaced along the experiment. The number of generations was generally similar among treatments, totalling 759, 815, and 830 generations for the parapatric, allopatric galactose, and sympatric treatments, respectively, except for the allopatric fructose treatment, which experienced 1603 generations (Supp. Fig. S1). Although two specialist phenotypes had emerged in all treatments at the final timepoint T3 (Fig. 2a), their trajectories differed among treatments (Fig. 3). In allopatry, fructose-patch strains slightly optimized growth on fructose, while remaining within the fructose-specialist phenotypic space (Fig. 3a, red). In the allopatric galactose patch, strains initially diversified into two phenotypes (T1, Fig. 3a, blue): a potential generalist improving growth on both sugars and a galactose specialist (GS_*A*_) trading off performance on fructose to increase growth on galactose. By T2 and T3, only GS_*A*_ persisted, indicating competitive exclusion of the generalist strains (Fig. 3a, blue). The absence of a generalist phenotype and the persistence of the fructose and galactose specialists - FS and GS - in the parapatric and, especially, the sympatric treatments further support that the evolution of a generalist is constrained (Fig. 3b,c). In parapatry, the polymorphism and emergence of GS_*P*_ was delayed until T2 (vs. T1 in the allopatry and sympatry, Fig. 3b), but then adaptation accelerated, with GS_*P*_ slightly surpassing GS_*A*_ in galactose growth (0.74 h^*−*1^ vs. 0.73 h^*−*1^, Fig 3a,b). In contrast, the sympatric treatment showed delayed but no accelerating adaptation, with GS_*S*_ reaching only 0.64 h^*−*1^ at T3 (Fig. 3c).

The temporal analysis of gene variants supports the idea that adaptation is delayed for GS in parapatry and both FS and GS in sympatry, compared to allopatry (Fig. 5). Population-level sequencing identified parallel evolution of FS- and GS-specific new variants across replicates and treatments, with most variants arising in allopatry and parapatry, and only a few variants in sympatry (Fig. 5a). Notably, the parapatric treatment features more variants in *fbp* and *ptnABCD* than the allopatric treatment. Although the number of evolved variants in parapatry increases over time, reaching numbers similar to allopatry, their frequencies remain lower than those in the allopatric treatment (Fig. 5b). The higher frequencies observed in FS in allopatry compared to FS in parapatry could be related to the larger number of generations of the fructose allopatric treatment (Supp Fig. S1), providing more time for mutations to spread throughout the population. In sympatry, FS variants were absent, and GS variants remained rare and at low frequency (Fig. 5). This pattern may partially reflect the smaller effective population size of the sympatric treatment (60 ml vs. 120 ml in the other treatments; Fig. 1b,c,d). In addition, GS variants in sympatry were sampled from a polymorphic population dominated by FS (75% FS vs. 25% GS; Fig. 3c, T3), whereas in allopatry and parapatry they were sampled from the galactose patches. The lack of FS-associated mutations suggests that FS strains remained close to the ancestral phenotype. Alternatively, FS strains in sympatry may not have evolved *pfk* and *fbp* mutations because they avoided the anabolic constraint of strict fructose consumption by co-metabolizing a small fraction of galactose. This treatment structure is consistent with the restricted recurrence analysis, which recovered the strongest enrichment when focal mutations were tested within the subsets of lineages in which the corresponding specialist phenotype was expected to evolve (Supp. Table S5).

**FIGURE 5.**
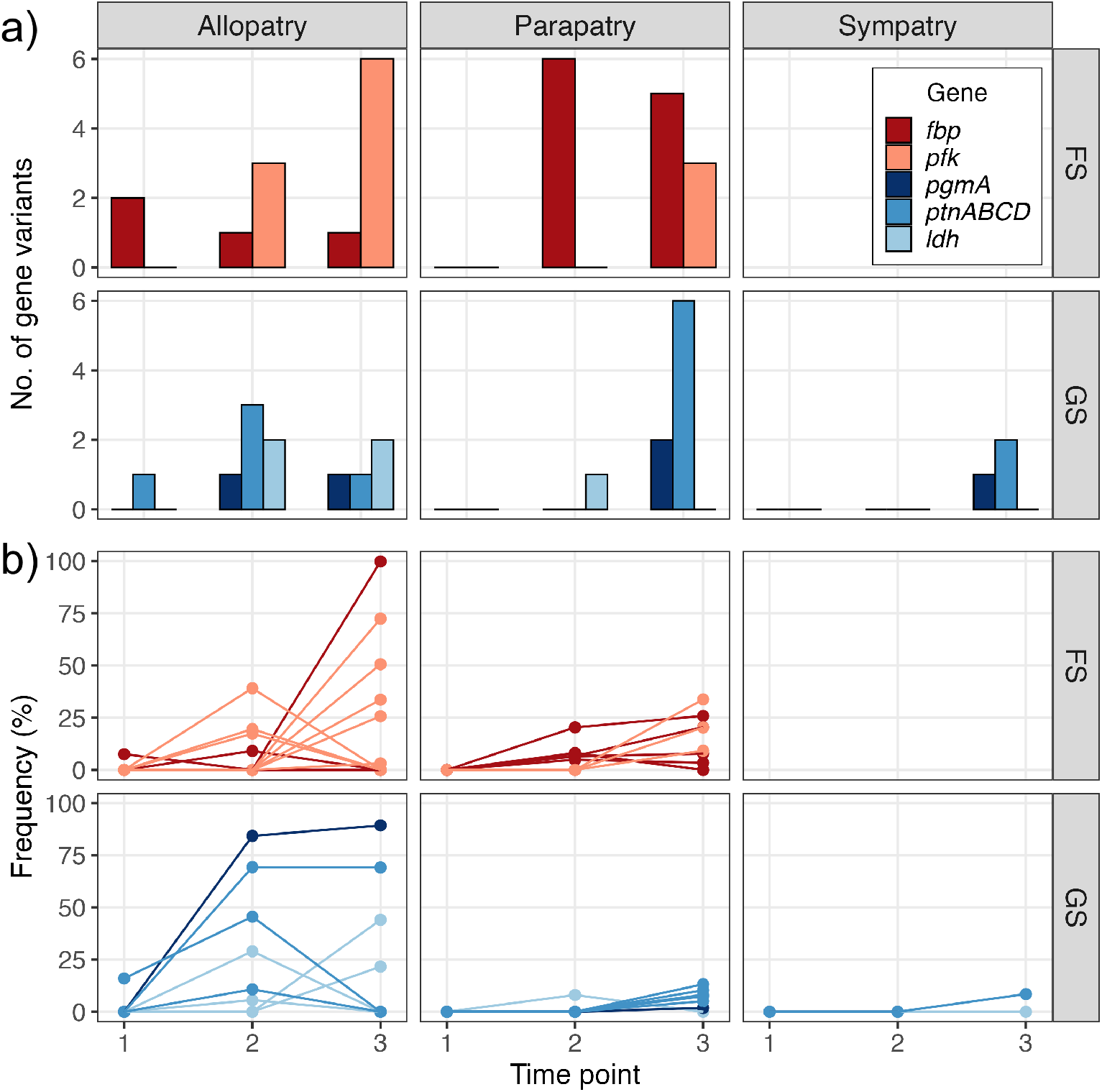
Variant counts and frequencies associated with fructose specialists (FS) and galactose specialists (GS) during the evolution experiment. **(a)** Total counts of evolved variants of FS (*fbp* and *pfk*) and GS (*pgmA, ptnABCD*, and *ldh*) loci by treatment and timepoint. **(b)** Frequency of variants through time. Note that not all SNP/indel nor CNV variants that were identified from the single strain analysis could be detected in the metagenome population samples.

### 3.5 Migration load and competitive ability in parapatry

The migration rate between the two parapatric patches was fixed for each patch at 5% flux of culture compared to dilution rate of the chemostat (Fig. 1c), regardless of local population density. Culture density, however, was consistently higher in the fructose than in the galactose patch (Fig. 6b), producing a density-asymmetry that imposed a stronger migration load from the fructose to the galactose patch than in the opposite direction (source-sink dynamics; Fig. 6a). By timepoint T3, the density difference between patches decreased due to the high relative growth improvements of the GS_*P*_ compared to FS_*P*_ (Fig. 6b, T3; see also Fig. 4b), resulting in a more balanced migration load between patches.

**FIGURE 6.**
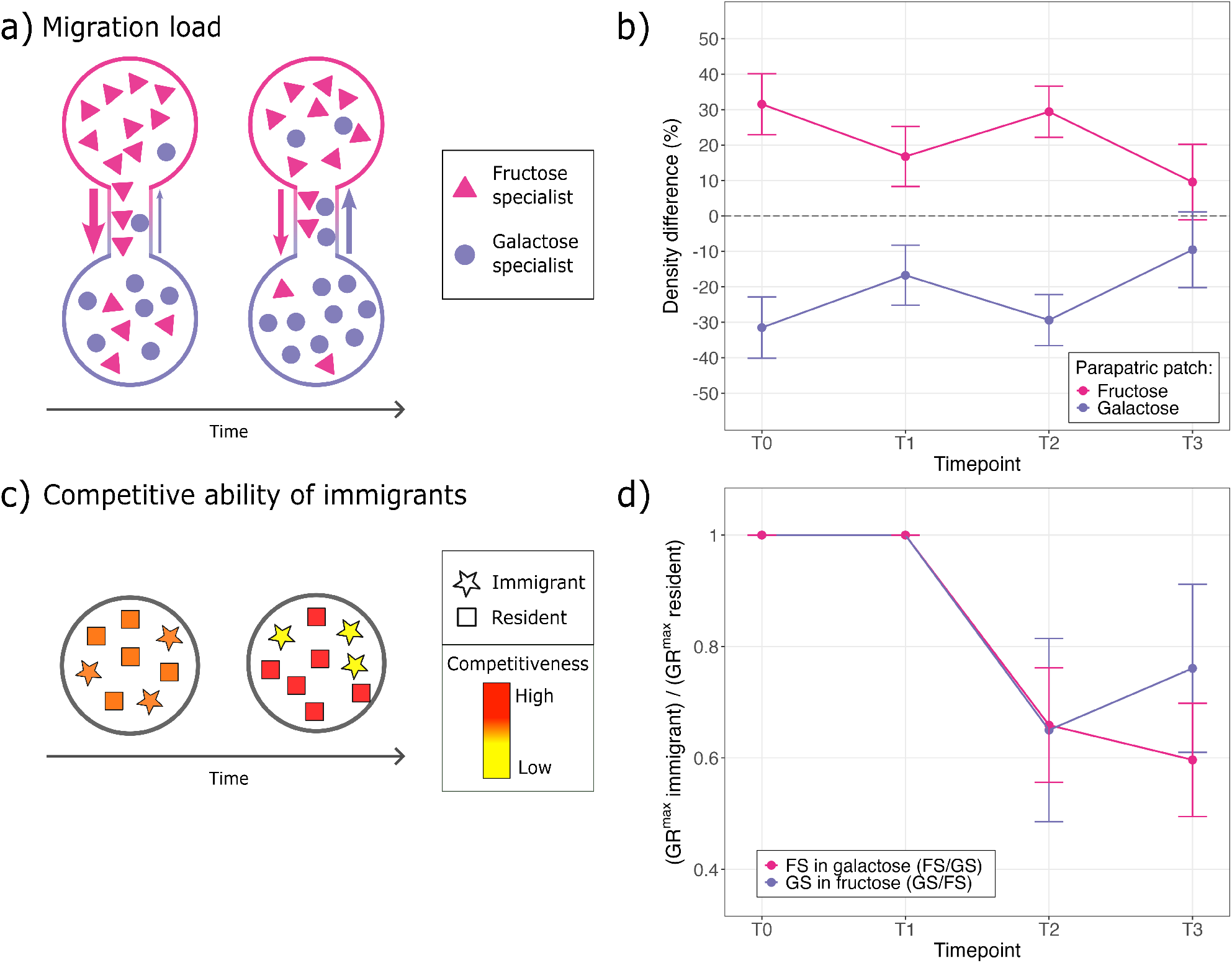
**(a)** Source-sink dynamics are expected to decrease through time because of increased growth rates and subsequent population density increase of the sink population. **(b)** Percent difference between the population density of the parapatric treatment (i.e., average between both patches, per replicate) and the density of each patch (per replicate) (fructose: FS_*P*_ and galactose: GS_*P*_). Population density was measured from the absorbance of chemostat cultures (OD_600_). **c)** The competitive ability of immigrants is expected to decrease over time because of local adaptation of residents. **(d)** Change of competitive ability of immigrants through time. The y-axis is the ratio between the immigrants’ and the residents’ maximum growth rates (GR^*max*^). For example, the fructose specialist (FS) is the immigrant in the galactose patch and the galactose specialist (GS) is its resident; thus the GR^*max*^ ratio is calculated as FS/GS and is above one if the FS does better than GS in galactose and below one if FS is worse than GS in galactose (as observed). The opposite is valid for the fructose patch. GR^*max*^ was derived from monocultures that were grown in CDMPC supplemented with either fructose or galactose.

We then investigated the within-patch dynamics in the parapatric treatment by quantifying the impact of immigrants on the local resident population (Fig. 6c). Immigrant competitive ability - their capacity to contribute to new generations relative to residents - was measured by comparing the growth rates of the local (resident) versus non-local (immigrant) sugar patch. Accordingly, we compared GS_*P*_ and FS_*P*_ growth rates (from the same replicate and timepoint) on both galactose and fructose sugars (Fig. 6d). In T1, immigrants and residents performed equally. In T2, immigrant competitive ability dropped sharply in both patches, creating an asymmetry in maximum growth rates between residents and immigrants within each patch. By T3, competitive ability decreased more slowly for FS_*P*_ immigrants and even showed a slight, non-significantly, increase for the GS_*P*_ immigrants. Overall, the immigrant’s competitive ability decreased over time as a result of residents’ local resource specialization through time.

In conclusion, the combined decrease in migration load (Fig. 6a,b) and immigrant competitive ability (Fig. 6c, d) through time led to a progressively smaller net impact of migration on the locally evolving populations. This pattern aligns with the delayed adaptation observed in parapatric evolutionary trajectories (Fig. 3).

## 4 DISCUSSION

### 4.1 Local resource availability and metabolic constraints are dominant drivers of spatial diversification

Heterogeneity in resource availability drove the evolution of resource specialists in our experimental populations of *Lactococcus cremoris*. Two distinct specialist phenotypes - fructose specialists (FS) and galactose specialists (GS) - evolved independently across all treatments, including under continuous gene flow in parapatry (Fig. 2a). These specialists optimized growth on one sugar at the cost of performance on the other, reflecting a strong metabolic trade-off. Genetic and transcriptomic analyses revealed that these specialists arose through mutations at key metabolic junctions where fructose and galactose pathways require opposite flux directions relative to the ancestral glucose metabolism (Fig. 4; Ekkers et al. 2022). These adaptations also induced consistent pleiotropic effects on growth across multiple carbon resources (Fig. 2b), suggesting that pathway structure constrains the evolutionary feasibility of different resource strategies and ultimately drives diversification.

Importantly, despite differences in spatial structures, similar phenotypes evolved across all three treatments, and these were often associated with mutations affecting the same genes or metabolic functions. The observed convergence indicates that local resource availability and underlying metabolic trade-offs were the primary factors determining the evolutionary outcomes. This conclusion is further supported by the evolution of a distinct trade-off shape observed in all treatments. In contrast to a gradual, continuous buildup of trade-off strength expected under mutation accumulation, the trade-off emerged around timepoints T1 and T2 and remained largely constant - or slightly weakened - by T3 (Fig. 3). Although many sequenced strains do not contain mutations in the identified targets of selection but do feature the phenotypic characteristics of the identified FS and GS specialists, we suspect that there are unannotated or uniquely evolved adaptive variants that were not identified by this study. The pertaining challenge in identifying function to poorly annotated genes, combined with labor-intensive functional testing of potentially impactful variants, limits the scope of the analysis of evolved variants.

Our findings highlight that trade-offs underlying local adaptation need not arise primarily through the accumulation of neutral mutations that become deleterious in alternative environments, as commonly observed in asexual microbial evolution experiments [Bono et al., 2017, Maclean, 2005], but can also emerge directly from antagonistic pleiotropy when the underlying metabolic architecture imposes strong functional constraints. Our results suggest that antagonistic pleiotropy, in which mutations beneficial for one resource simultaneously reduce performance on others, can also drive local adaptation. Whether divergence among patches is driven primarily by antagonistic pleiotropy or by mutation accumulation likely depends on the strength and shape of trade-offs in combination with gene flow rates. The formal recurrence analyses support this interpretation, but also clarify its limits. Across the population-level sequencing data, recurrent gene hits occurred more often than expected by chance, supporting the inference that selection repeatedly targeted shared loci (Supp. Table S4). At the same time, the focal mutations identified from the phenotypic and functional analyses were not expected to recur uniformly across all 20 lineages. Instead, the clearest enrichment was recovered when these targets were tested within ecologically relevant subsets of lineages. In this analysis, pfk showed strong enrichment in fructose-specialist lineages, whereas fbp and ldh followed the expected directional pattern with weaker support, and pgmA and ptnABCD remained too sparse in the pooled population data for strong formal significance (Supp. Table S5).

Our study reveals that understanding the relation between functions under selection is crucial for predicting evolutionary outcomes. Migration between patches did not prevent diversification: populations consistently evolved into a polymorphism of two sugar specialists (Fig. 2a). These results demonstrate that metabolic constraints, together with local resource heterogeneity, can promote diversification even in the presence of gene flow [Ravigné et al., 2009].

### 4.2 Migration load and competitive dynamics create eco-evolutionary feedbacks that determine the tempo of diversification

Despite the similarities among the evolved phenotypes across treatments (Fig. 2a), the tempo of divergence differed markedly. In the parapatric treatment, phenotypic adaptation was delayed relative to the allopatric control, with divergence emerging only at T2 rather than T1 (Fig 3). This lag is also observed in the number and frequency of gene variants (Fig. 5). However, adaptation then accelerated, ultimately matching or exceeding allopatric adaptation by T3. This biphasic temporal pattern reveals dynamic eco-evolutionary feedback between migration and local adaptation.

This initial delay can be explained by source-sink dynamics driven by asymmetric habitat suitability. The ancestor strain was optimally adapted to glucose, which biased resource use toward fructose (Fig. 1a). Consequently, in the early stage of the experiment, population density was higher in the fructose patch than in the galactose patch (Fig. 6a,b), leading to substantial gene flow from fructose to galactose. Because the migration rate was fixed at 5% regardless of local density, this asymmetry imposed a disproportionate migration load on the galactose patch, slowing the establishment of galactose specialists. Indeed, initial differences in habitat suitability can result in differences in population density between habitats and strong source-sink dynamics [Bisschop et al., 2019]. Such differences in population density among habitats are indicative of hard selection, which is less conducive to promoting diversification [Ravigné et al., 2009].

As galactose specialists evolved and improved growth on galactose, however, two reinforcing changes occurred. First, population density in the galactose patch increased, reducing the density asymmetry between patches (Fig. 6b). This demographic shift lowered the relative migration load from fructose to galactose. Second, and critically, the competitive ability of immigrants declined as residents specialized (Fig. 6d). By T2 and T3, immigrants performed substantially worse than residents on local resources, reflecting the metabolic trade-offs imposed by specialization. Together, these eco-evolutionary feedbacks reduced the effective migration rate over time, progressively weakening migration’s constraining effect and allowing accelerated local adaptation. This interpretation is further supported by the temporal pattern of mutation frequencies: galactose-specialist variants increased more slowly in parapatry than allopatry early on but reached comparable levels by T3 (Fig. 5b). Importantly, these feedbacks were not merely ecological responses or evolutionary responses in isolation, but emerged from their dynamic interaction: ecological conditions (density asymmetries, competition) constrained evolution early on, but evolutionary changes (local adaptation, specialization) fed back to alter ecological conditions, ultimately facilitating further divergence.

Our findings help reconcile apparently contradictory results from previous spatial evolution experiments. Two recent yeast studies found that intermediate gene flow (parapatry) constrained adaptive divergence more than either complete isolation (allopatry) or full mixing (sympatry) [Tusso et al., 2021, Gray and Goddard, 2012a]. These studies concluded that parapatric gene flow inhibits local adaptation. One possible explanation for the difference with our results is that those experiments were shorter in duration and may have captured an earlier phase of adaptation, in which migration load strongly constrains local divergence. However, an additional difference is that both yeast systems involved recurrent sexual reproduction and recombination. In such systems, recombination can facilitate adaptation by combining beneficial alleles across backgrounds and reducing selective interference, but it can also disrupt favorable allele combinations. By contrast, in our asexual system, adaptation depends entirely on the sequential accumulation of linked mutations. Reproductive mode may therefore have contributed to the contrasting outcomes, together with other differences between systems, including the availability of standing variation and the genetic architecture of the selected traits.

### 4.3 Effects of differences in effective population size

An independent factor that may have contributed to the observed delay and acceleration of local adaptation in the parapatric treatment relates to the availability of a larger gene pool and parallel selection pressures in a spatially heterogeneous environment with multiple subpopulations. All else being equal, the experimental setup of the parapatric treatment allows for a population size twice as large as in the sympatric treatment (because it is composed of two connected chemostats, Fig. 1b,c,d) and for gene exchange between patches as opposed to the allopatric treatment. Thus, adaptations requiring a combination of multiple mutations, such as reciprocal sign epistasis, might evolve more easily in this larger, multi-patch environment. These epistatic adaptations could have postponed the need to trade off fructose and galactose metabolism, thereby delaying divergence of the local specialists. We attempted to test this hypothesis by means of phylogenetic comparison of specialists from the different treatments. However, the high proportion of strain-specific mutations (Supp. Fig. S7) prevented a reliable reconstruction of phylogenetic trees, limiting our ability to track the ancestry of the adaptive strains within the parapatric patches. In addition, differences in the number of generations among treatments may have affected mutational opportunities for FS, highlighted by higher rates of fixation of evolutionary targets (Fig. 5a). However, these differences do not appear to explain the main treatment-level patterns related to moderate FS growth rate improvements on fructose (Fig. 4a,b) and a delayed divergence of GS phenotypes in the parapatric treatment, where there was no comparable outlier in the number of generations nor population density.

In contrast to our asexual system, sexual populations can exchange beneficial alleles across lineages through recombination, which may reduce selective interference and facilitate adaptation by both combining beneficial mutations and unlinking beneficial from deleterious mutations [Gray and Goddard, 2012b, Paree and Teotonio, 2025, Livnat et al., 2008, Keightley and Eyre-Walker, 2000, Otto and Gerstein, 2006]. However, recombination can also break up favorable allele combinations and thereby alter the tempo and genetic basis of divergence. In our system, the high number of parallel adaptive variants within the same replicate at relatively low frequency in the parapatric treatment (Fig. 5) suggests that populations were constrained by clonal interference and that horizontal gene transfer did not readily occur.

### 4.4 Broader implications for ecological and evolutionary theory

Our results demonstrate that predicting spatial divergence requires not only an understanding of migration rates or trade-off strength in isolation, but also their dynamic interaction mediated by ecological feedbacks. Theoretical models of adaptive diversification typically treat migration as a static parameter opposing local adaptation [Doebeli and Dieckmann, 2003]. Our findings suggest that this framework is incomplete: the effect of migration on divergence changes as populations adapt, creating eco-evolutionary feedback loops: evolutionary changes modify the ecological conditions that, in turn, shape further evolution.

This eco-evolutionary perspective has important implications. First, it suggests that short-term observations, whether from experiments or field studies, may be misleading for predicting long-term outcomes. Systems that initially appear constrained by gene flow may eventually diversify as eco-evolutionary feedbacks alter the ecological conditions. Second, it highlights that preadaptation and initial asymmetries critically influence whether and when divergence occurs. Third, it emphasizes that the same migration rate can have different effects depending on the eco-evolutionary context: high migration may either prevent or facilitate divergence depending on density regulation, competitive interactions, and the stage of adaptation.

In conclusion, spatial diversification in heterogeneous environments arises from the interplay between resource-utilization trade-offs, migration load, and competitive dynamics. In our experiment, selection for efficient growth on alternative resources subject to metabolic constraints consistently drove specialization regardless of spatial structure, yet eco-evolutionary feedbacks between population density, competitive ability, and migration load shaped the tempo of divergence. These results demonstrate that ecological conditions and evolutionary processes form feedback loops that dynamically govern both the rate and outcome of adaptive diversification. Understanding this feedback is essential for predicting when and how spatial heterogeneity promotes diversity in nature.

## AUTHOR CONTRIBUTIONS

D.M.E. conceived and designed the study, performed the experiments, phenotypic characterization and genome extractions, led the data analyses, and wrote the manuscript. S.T. performed the individual and population sequence analyses and edited the manuscript. M.C.R. performed the clustering analysis, co-created the figures, and edited the manuscript. O.P.K. provided laboratory support, and experimental and genetic expertise for the conception of the project. G.S.v.D. co-conceived the idea, provided advice throughout the experiment, and edited the manuscript. All authors contributed to the interpretation of the data and commented on the manuscript.

## ACKNOWLEDGMENTS

We thank Stefany Moreno for helping with the analysis of growth rate and yield data from the growth curves; Cyrus Mallon for help with running the evolution experiment; Anne-Marie Veenstra-Skirl for assisting with the plating experiments; Anne de Jong for advising on the sequencing data analysis; Harma Karsens for performing the RNA extractions and qPCR experiments. This work was supported by the European Research Council (ERC Starting Independent Researcher Grant 309555 to G.S.v.D.) and the Netherlands Organization for Scientific Research (NWO Vidi Grant 864.11.012 to G.S.v.D.).

## ETHICAL STATEMENT

This study did not involve human participants or animals and therefore did not require ethical approval.

## DATA AVAILABILITY STATEMENT

The data are not publicly available online but can be shared upon request by contacting the corresponding author.

