## Supplementary material for "Migration load, competition, and metabolic trade-offs shape spatial divergence through eco-evolutionary dynamics"

602 SUPPORTING INFORMATION

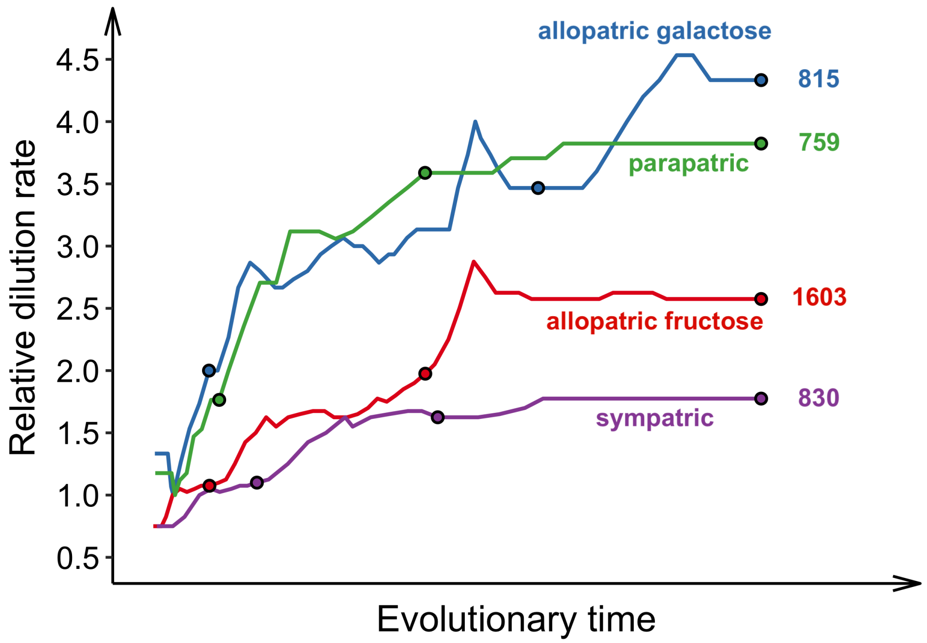


**FIGURE S1** Relative daily dilution rates throughout the evolution experiment by treatment. Relative dilution rate, used here as an approximation of relative growth-rate improvement, was calculated as the dilution rate of evolved populations divided by the maximum dilution rate of the ancestral strain in the corresponding resource environment. The number of generations experienced by each treatment at the end of the evolution experiment is indicated as a number on the right side of each line. The dots indicate the three timepoints sampled for each treatment. For more details on the experimental procedures and the calculation of dilution rates and number of generations, see Methods section 2.1.

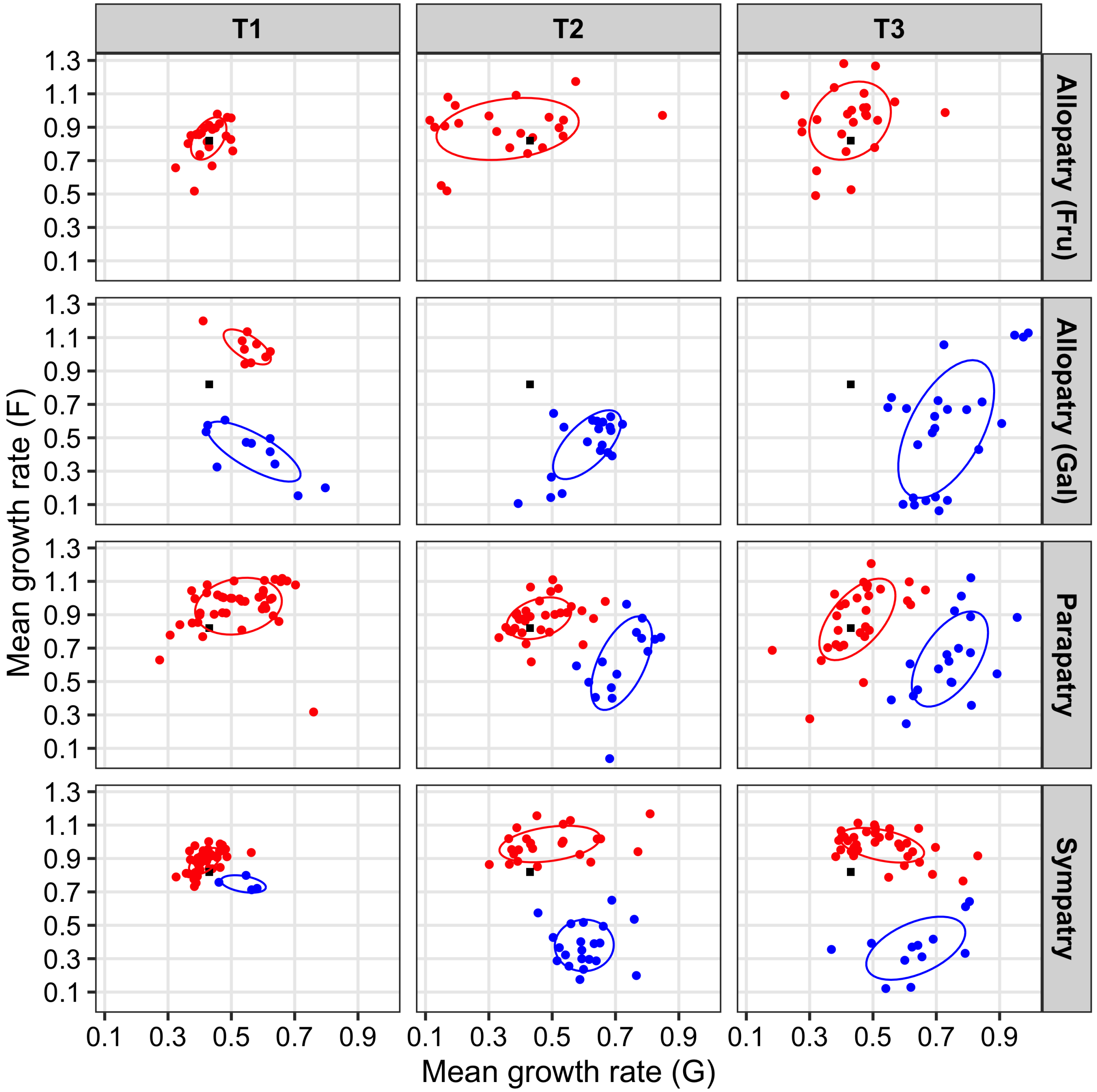


**FIGURE S2** Phenotypic clustering analysis. Clustering was performed on the maximum growth rate coordinates on fructose (F) and galactose (G) for each of the genotypes sampled from the evolving populations across the treatments (rows) at timepoint T1, T2 and T3 (columns). The two colors indicate the different phenotypic clusters.

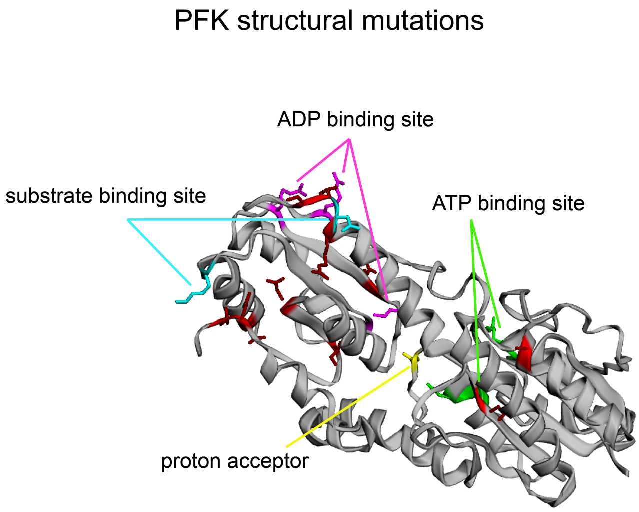


**FIGURE S3** Pfk structural mutations. Mutations (red) mapped to Pfk protein and its ATP binding site (green), ADP binding sites (purple), substrate binding site (turquoise) and proton acceptor (yellow). Figure generated using EzMol [Reynolds et al., 2018].

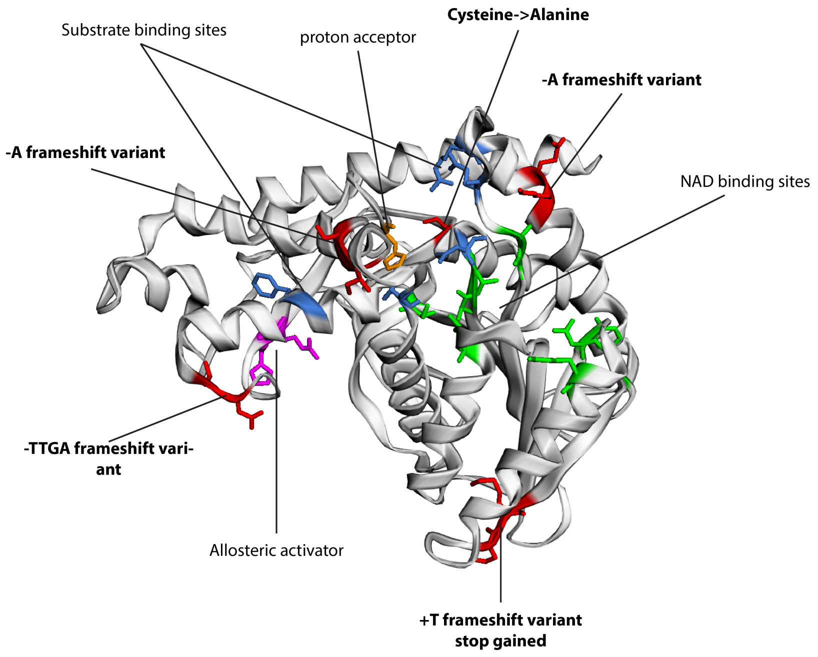


**FIGURE S4** Mutations (red) mapped to LDH protein and its substrate binding site (blue), NAD+ binding site (green), allosteric activator site (purple), and proton acceptor (yellow). Figure generated using EzMol [Reynolds et al., 2018].

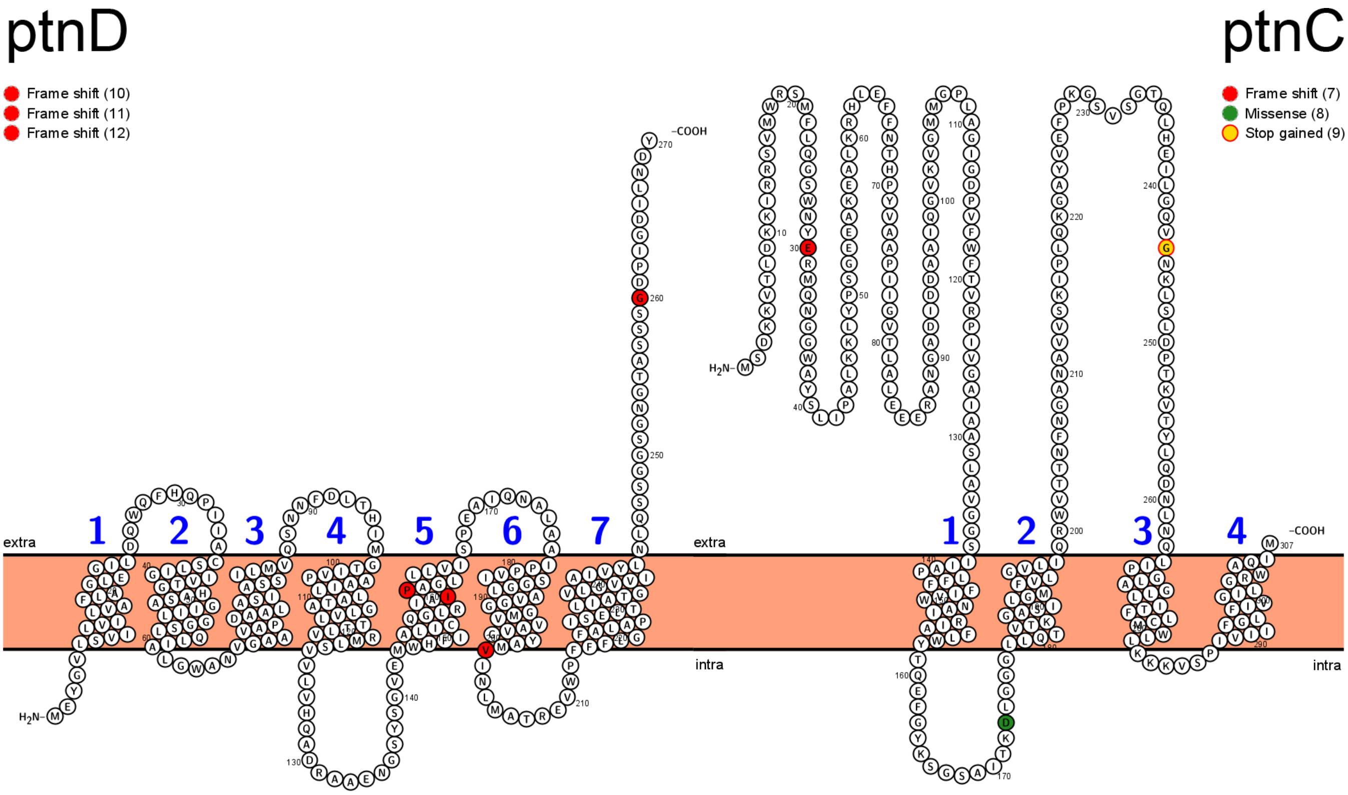


**FIGURE S5** Structural mutations mapped to ptnC and ptnD transmembrane protein elements. See figure legend for types of mutations. Figure generated using Protter [Omasits et al., 2014].

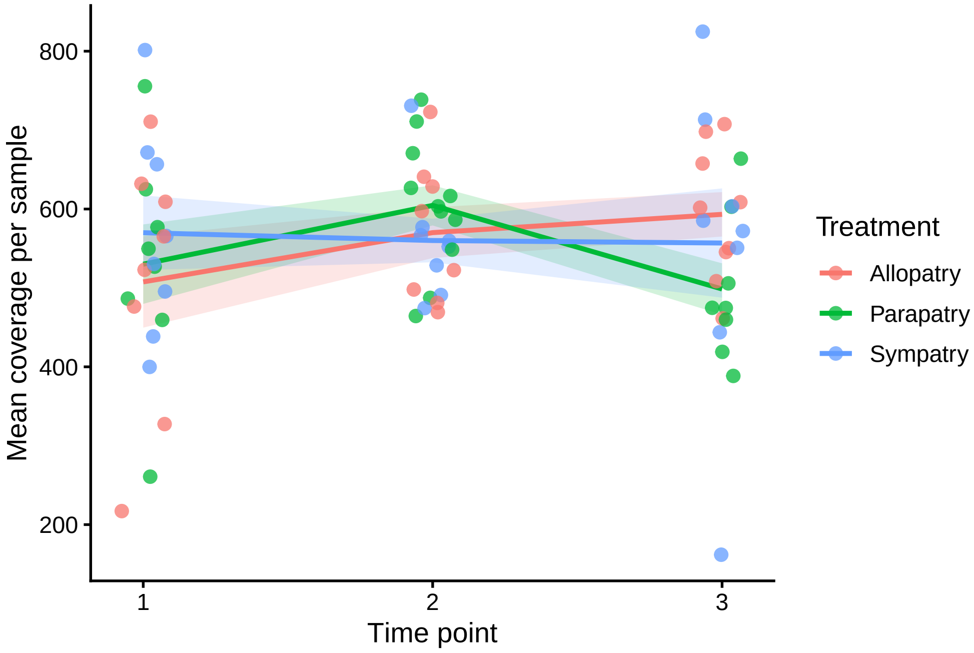


**FIGURE S6** Mean whole-genome coverage per sequenced population sample across timepoints and treatments. Points show mean coverage for individual samples, and colored lines indicate treatment-level trends across time-points. Across all sequenced samples, the average coverage was 557×, with a minimum of 162× and a maximum of 825×.

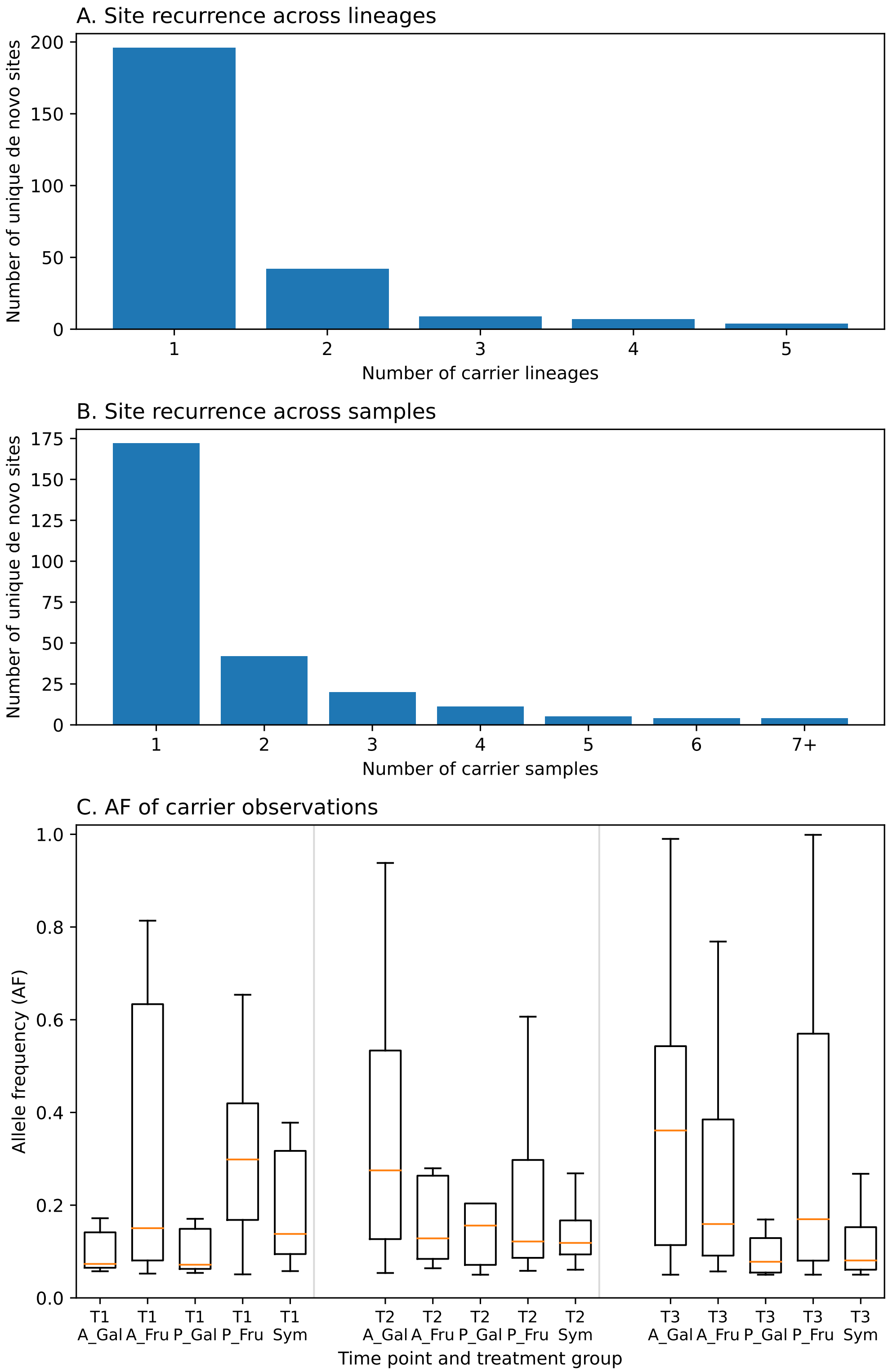


**FIGURE S7** Genome-wide summary of de-novo mutations detected in the population-level sequencing data across all timepoints after filtering against the ancestral strain. (a) Distribution of unique de-novo sites by the number of independent carrier lineages. (b) Distribution of unique de-novo sites by the number of carrier population samples. (c) Allele-frequency distributions of carrier observations by timepoint and treatment group. In panel (c), A_Gal = allopatric galactose, A_Fru = allopatric fructose, P_Gal = parapatric galactose patch, P_Fru = parapatric fructose patch, and Sym = sympatry. Most de-novo sites were lineage-specific (196/258), and only 62/258 recurred across more than one lineage.

**TABLE S1** List of selected strains (and their respective mutations) used for functional tests (expression and enzymatic).

| Strain name | Phenotype | Treatment Occurence | Replicate | Timepoint | Mutations |
|---|---|---|---|---|---|
| Anc. | Ancestor | | | 0 | None |
| $FS_{a1}$ | Fructose specialist | Allopatry (Fru) | 4 | 1 | *fbp* CNV220201 two-fold duplication of 81498 bp section including *fbp* |
| $FS_{a2}$ | Fructose specialist | Allopatry (Fru) | 3 | 3 | *fbp* (intergentic SNP, C-250825→A), *pfk C1078234→A* (missense SNP, alanine18→glutamic acid) |
| $FS_{p1}$ | Fructose specialist | Parapatry | 4 | 3 | population sample (*pfk* missense variant A1078681 >G lysine>arginine, *fbp* intergenic SNP T250792→C) |
| $FS_{p2}$ | Fructose specialist | Parapatry | 3 | 3 | *fbp* (intergenic SNP, G250865→CA) |
| $FS_{s1}$ | Fructose specialist | Sympatry | 4 | 3 | no mutations in *fbp* or *pfk* |
| $GS_{a1}$ | Galactose specialist | Allopatry (Gal) | 2 | 3 | *pgmA* (intergentic SNP, T540043→A), *ptnABCD* (intergentic insetion A712481→ATCGATATTTG), *ldh* (frameshift variation CTA1081096→CTTA) |
| $GS_{a2}$ | Galactose specialist | Allopatry (Gal) | 2 | 3 | *pgmA* (intergentic SNP, T540043→A), ptnABCD (intergentic insetion A712481→ATCGATATTTG) |
| $GS_{p1}$ | Galactose specialist | Parapatry | 3 | 3 | *ptnAB* ( deletion intergenic GTTATTATAC712464→GTTATAC) , |
| $GS_{p2}$ | Galactose specialist | Parapatry | 3 | 2 | *ldh* (missense variant T1081725→C) |
| $GS_{s1}$ | Galactose specialist | Sympatry | 4 | 3 | *pgmA* CNV536201 six-fold duplication of 8699 bp section including *pgmA* |

**TABLE** S2 Primer sequences used in the qRT-PCR expression experiments to measure expression levels of mutated genes (*fbp*, *pfk*, *pgmA* and *ptnAB*) and housekeeping gene (*glyA*) of ancestor and mutants strains.

| Gene | Primer |
|---|---|
| glyA_FW | ATGGGAAAGAAGCACAGGACTTATTAG |
| glyA_RV | TGCCGCACCAATTCGCAC |
| fbp_FW | TCGCAACTACAAAATCTGGGAGAAC |
| fbp_RV | CGCGGTCATAAATATCCCCAACAAC |
| pfk_FW | ACATTCTTGTACTCAGCACGTTACC |
| pfk_RV | CACCACCGATTACAACGACACC |
| pgmA_FW | TGTTGCTCAAGGAACTCAATATTATGCTC |
| pgmA_RV | AGCTGCTTTTTCAAGTGCTCCC |
| ptnAB_FW | AAATTGAAGCTGCCATCGCAAC |
| ptnAB_RV | TGGATTTTCACCCATCACTGCAC |

**TABLE S3** List of evolved unique mutation variants identified from monoculture strains and population-level sequencing during the experiment. The column *Occurrence* indicates whether each variant was detected in monoculture samples (Mono), population samples (Pop.), or both. The table includes a strain-specific identifier (*Strain IDs*) to indicate which variants occurred in the same sequenced strain; and reports variant frequency where this could be estimated from the population sequencing data at the sampled time points (*fqT1*, *fqT2*, *fqT3*). Variants detected only in the single-strain analysis do not have a population-frequency estimate. Note that some variants were detected at consecutive time points.

| Gene | Variant type | Mutation type | Position | mutational change | Variant nr. | Occurence | Strain IDs | Treatment | Timepoint | Amino acid change | fq T1 | fq T2 | fq T3 |
|---|---|---|---|---|---|---|---|---|---|---|---|---|---|
| Fructose specialist | | | | | | | | | | | | | |
| fbp | cnv | duplication (2x) | 220201 | 81498 | 1 | Mono (1x) | 1 | Allopatric | 1 | | NA | | |
| fbp | snps | intergenic_region | 250787 | G -> A | 2 | Pop. | | Parapatric | 3 | | | 4.28 | |
| fbp | snps | intergenic_region | 250787 | G -> A | 2 | Pop. | | Parapatric | 2.3 | | | 20.38 | 25.87 |
| fbp | snps | intergenic_region | 250791 | G -> A | 3 | Pop. | | Parapatric | 2 | | | 7.81 | |
| fbp | snps | intergenic_region | 250791 | G -> A | 3 | Pop. | | Parapatric | 2.3 | | | 6.63 | 20.49 |
| fbp | snps | intergenic_region | 250791 | G -> A | 3 | Pop. | | Parapatric | 3 | | | | 3.03 |
| fbp | snps | intergenic_region | 250792 | C -> T | 4 | Pop. | | Parapatric | 2.3 | | | 4.9 | 3.43 |
| fbp | snps | intergenic_region | 250825 | C -> T | 4 | Pop. + Mono (3x) | 2;3;4 | Allopatric | 1.3 | | 7.57 | | 99.82 |
| fbp | snps | intergenic_region | 250825 | C -> T | 4 | Pop. + Mono (2x) | 5;6 | Allopatric | 2 | | | 9.09 | |
| fbp | snps | intergenic_region | 250825 | C -> T | 4 | Pop.+ Mono (1x) | 7 | Parapatric | 2 | | | 8.12 | |
| fbp | snps | intergenic_region | 250865 | GCAG -> GG | 5 | Mono (1x) | 8 | Parapatric | 3 | | | | NA |
| fbp | del | intergenic_region | 250865 | G -> CA | 6 | Pop. | | Parapatric | 2.3 | | | 6.48 | 7.84 |
| pfk | snps | missense_variant | 1078234 | C -> A | 1 | Pop. + Mono (2x) | 4;5 | Allopatric | 3 | alanine --> glutamic acid | | | 20.32 |
| pfk | snps | missense_variant | 1078234 | C -> A | 1 | Pop. + Mono (2x) | 6;7 | Allopatric | 2 | alanine --> glutamic acid | | 19.57 | |
| pfk | snps | missense_variant | 1078545 | C -> T | 2 | Pop. | | Parapatric | 3 | leucine -> phenylalanine | | | 9.21 |
| pfk | snps | missense_variant | 1078641 | C -> T | 3 | Pop. | | Allopatric | 3 | arginine -> cysteine | | | 72.41 |
| pfk | snps | missense_variant | 1078648 | C -> T | 4 | Pop. | | Allopatric | 2 | threonine -> isoleucine | | 39.13 | |
| pfk | snps | missense_variant | 1078651 | C -> T | 5 | Pop. | | Allopatric | 3 | serine -> phenylalanine | | | 33.67 |
| pfk | snps | missense_variant | 1078651 | C -> T | 5 | Pop. + Mono (2x) | 10;11 | Allopatric | 2 | serine -> phenylalanine | | 17.36 | |
| pfk | snps | missense_variant | 1078681 | A -> G | 6 | Pop. | | Parapatric | 3 | lysine -> arginine | | | 33.81 |
| pfk | snps | missense_variant | 1078741 | A -> C | 7 | Pop. | | Allopatric | 3 | asparic acid -> arginine | | | 25.73 |
| pfk | snps | missense_variant | 1078753 | T -> C | 8 | Pop. | | Allopatric | 3 | methionine -> threonine | | | 3.16 |
| pfk | snps | missense_variant | 1078780 | A -> G | 9 | Pop. | | Allopatric | 3 | asparagine -> serine | | | 50.6 |
| pfk | indels | inframe deletion | 1079189 | TCTTAACCTTAACT -> TCTTAACT | 10 | Mono (1x) | 12 | Allopatric | 3 | leucine and aspagine deleted | | | NA |
| Galactose specialist | | | | | | | | | | | | | |
| ptnABC | cnv | deletion (0,5x) | 706001 | 10698 | 1 | Pop. | | Sympatric | 3 | | 49 | | |
| ptnD | in | frameshift_variant | 710068 | A -> TT | 2 | Pop. + Mono (1x) | 19 | Parapatric | 3 | | | | 5.1 |
| ptnD | in | frameshift_variant | 710199 | A -> AC | 3 | Pop. | | Parapatric | 3 | | | | 8.17 |
| ptnD | in | frameshift_variant | 710375 | A -> AG | 4 | Pop. | | Parapatric | 3 | | | | 10.34 |
| ptnC | in | frameshift_variant | 710631 | G -> GA | 5 | Pop. | | Parapatric | 3 | | | | 7.4 |
| ptnC | snps | missense_variant | 711058 | C -> T | 6 | Pop. | | Allopatric | 1.2 | alanine -> valine | 16 | 45.57 | |
| ptnC | snps | stop gained | 711271 | C -> T | 7 | Pop. | | Parapatric | 3 | | | | 13.2 |
| ptnAB | snps | intergenic_region | 712439 | A -> G | 8 | Pop. | | Sympatric | 3 | | | | 8.5 |
| ptnAB | del | intergenic_region | 712464 | GTTATTATAC -> GTTATAC | 9 | Mono (1x) | 20 | Parapatric | 3 | | | | NA |
| ptnAB | snps | intergenic_region | 712468 | T -> C | 10 | Pop. | | Allopatric | 2 | | | | 10.75 |
| ptnAB | in | intergenic_region | 712481 | A -> ATCGATATTTGC | 11 | Pop. + Mono (5x) | 14;15;16;17;18 | Allopatric | 2.3 | | | 69.27 | 69.09 |
| pgmA | cnv | duplication (2x) | 528301 | 21098 | 1 | Pop. | | Parapatric | 3 | | NA | | |
| pgmA | cnv | duplication (6x) | 536201 | 8699 | 2 | Mono (1x) | 13 | Sympatric | 3 | | NA | | |
| pgmA | indels | intergenic_region | 540043 | T -> A | 3 | Pop. | | Parapatric | 3 | | | | 1.87 |
| pgmA | indels | intergenic_region | 540043 | T -> A | 3 | Pop. + Mono (5x) | 14;15;16;17;18 | Allopatric | 2.3 | | | 84.21 | 89.26 |
| ldh | in | stop gained | 1081096 | CTA -> CTTA | 1 | Pop. + Mono (1x) | 17 | Allopatric | 3 | | | | 44.02 |
| ldh | indels | frameshift_variant | 1081203 | G -> A | 2 | Pop. | | Allopatric | 2 | | | 5.61 | |
| ldh | indels | frameshift_variant | 1081410 | G -> TTGA | 3 | Pop. | | Allopatric | 3 | | | | 21.63 |
| ldh | indels | frameshift_variant | 1081588 | C -> A | 4 | Pop. | | Allopatric | 2 | | | 28.96 | |
| ldh | snps | missense_variant | 1081725 | T -> C | 5 | Pop. + Mono (1x) | 21 | Parapatric | 2 | cysteine -> alanine | | 8.07 | |

**TABLE S4** Genome-wide recurrence statistics for de novo gene hits in population-level sequencing data. Column *dataset* indicates whether the analysis was performed on the *T3*endpoint data or the full time-series dataset (*alltp*); *row_type* distinguishes genome-wide summary statistics (*global_summary*) from gene-level recurrence results (*gene_parallelism*); *statistic_or_gene* gives either the global statistic name or the gene identifier; *locus_tag*, *gene_name*, *feature_for_length*, and *target_length_bp* describe the annotated gene and the mutational target size used in the permutation test; *observed_hits* give the observed lineage occupancy, *null_mean_hits* and *null_sd_hits* give the mean and standard deviation of lineage occupancy under the permutation null model; *empirical_p_ge_obs* gives the empirical P-value; *q_bh* the Benjamini-Hochberg corrected q-value; and *lineages_hit* the lineages carrying *de-novo* mutations in the gene. **Supp. Table S4 available at https://doi.org/10.5281/zenodo.22051309**

**TABLE** S5 Restricted enrichment tests for focal specialist-associated targets. Column *dataset* indicates whether the analysis was performed on the *T3* endpoint data or the full time-series dataset (*alltp*); *target_label* gives the focal target used in the manuscript; *target_type* indicates whether it was tested as a coding gene or as a intergenic region; *source_id* gives the underlying gene or region identifier used in the analysis; *hypothesis_set* indicates whether the target was tested under the fructose-specialist (FS) or galactose-specialist (GS) expectation; *tested_subset* gives the ecologically relevant subset of lineages used for the restricted test: *FS_primary* = *allopatric-fructose* plus *parapatric-fructose* lineages, *FS_expanded* = *FS_primary* plus *sympatric lineages*, *GS_primary* = *allopatric-galactose* plus *parapatric-galactose* lineages, *GS_expanded* = *GS_primary* plus *sympatric* lineages; *n_total_lineages* gives the total number of independent lineages; *n_carrier_lineages_total* the total number of carrier lineages for the focal target; *n_subset_lineages* the number of lineages in the tested ecological subset; *n_carriers_in_subset* the number of carrier lineages within that subset; *carrier_lineages_total* the full set of carrier lineages; *subset_lineages* the lineages included in the tested subset; *p_enrichment* the one-sided hypergeometric P-value; and *q_bh* the Benjamini-Hochberg corrected q-value.

| dataset | target label | target type | source_id | hypothesis set | tested_subset | n_total_lineages | n_carrier_lineages_total | n_subset_lineages | n_carriers_in_subset | carrier_lineages_total | subset_lineages | p_enrichment | q_bh |
|---|---|---|---|---|---|---|---|---|---|---|---|---|---|
| T3 | pfk | gene | pfkA | FS | FS_primary | 20 | 5 | 8 | 5 | c10,c12,c18,c19,c20 | c09,c10,c11,c12,c17,c18,c19,c20 | 0.003611971 | 0.036119711 |
| T3 | pfk | gene | pfkA | FS | FS_expanded | 20 | 5 | 12 | 5 | c10,c12,c18,c19,c20 | c09,c10,c11,c12,c17,c18,c19,c20, c21,c22,c23,c24 | 0.051083591 | 0.255417957 |
| T3 | fbp | region | fbp | FS | FS_primary | 20 | 2 | 8 | 2 | c11,c19 | c09,c10,c11,c12,c17,c18,c19,c20 | 0.147368421 | 0.368421053 |
| T3 | fbp | region | fbp | FS | FS_expanded | 20 | 2 | 12 | 2 | c11,c19 | c09,c10,c11,c12,c17,c18,c19,c20, c21,c22,c23,c24 | 0.347368421 | 0.5 |
| T3 | ldh | gene | WP_003131075.1 | GS | GS_primary | 20 | 2 | 8 | 2 | c06,c07 | c05,c06,c07,c08,c13,c14,c15,c16 | 0.147368421 | 0.368421053 |
| T3 | ldh | gene | WP_003131075.1 | GS | GS_expanded | 20 | 2 | 12 | 2 | c06,c07 | c05,c06,c07,c08,c13,c14,c15,c16, c21,c22,c23,c24 | 0.347368421 | 0.5 |
| T3 | pgmA | region | pgmA | GS | GS_primary | 20 | 1 | 8 | 1 | c06 | c05,c06,c07,c08,c13,c14,c15,c16 | 0.4 | 0.5 |
| T3 | pgmA | region | pgmA | GS | GS_expanded | 20 | 1 | 12 | 1 | c06 | c05,c06,c07,c08,c13,c14,c15,c16, c21,c22,c23,c24 | 0.6 | 0.6 |
| T3 | ptnABCD | region | ptnABCD | GS | GS_primary | 20 | 1 | 8 | 1 | c06 | c05,c06,c07,c08,c13,c14,c15,c16 | 0.4 | 0.5 |
| T3 | ptnABCD | region | ptnABCD | GS | GS_expanded | 20 | 1 | 12 | 1 | c06 | c05,c06,c07,c08,c13,c14,c15,c16, c21,c22,c23,c24 | 0.6 | 0.6 |
| alltp | pfk | gene | pfkA | FS | FS_primary | 20 | 5 | 8 | 5 | c10,c12,c18,c19,c20 | c09,c10,c11,c12,c17,c18,c19,c20 | 0.003611971 | 0.036119711 |
| alltp | pfk | gene | pfkA | FS | FS_expanded | 20 | 5 | 12 | 5 | c10,c12,c18,c19,c20 | c09,c10,c11,c12,c17,c18,c19,c20, c21,c22,c23,c24 | 0.051083591 | 0.127708978 |
| alltp | fbp | region | fbp | FS | FS_primary | 20 | 3 | 8 | 3 | c11,c19,c20 | c09,c10,c11,c12,c17,c18,c19,c20 | 0.049122807 | 0.127708978 |
| alltp | fbp | region | fbp | FS | FS_expanded | 20 | 3 | 12 | 3 | c11,c19,c20 | c09,c10,c11,c12,c17,c18,c19,c20, c21,c22,c23,c24 | 0.192982456 | 0.275689223 |
| alltp | ldh | gene | WP_003131075.1 | GS | GS_primary | 20 | 3 | 8 | 3 | c06,c07,c15 | c05,c06,c07,c08,c13,c14,c15,c16 | 0.049122807 | 0.127708978 |
| alltp | ldh | gene | WP_003131075.1 | GS | GS_expanded | 20 | 3 | 12 | 3 | c06,c07,c15 | c05,c06,c07,c08,c13,c14,c15,c16, c21,c22,c23,c24 | 0.192982456 | 0.275689223 |
| alltp | pgmA | region | pgmA | GS | GS_primary | 20 | 1 | 8 | 1 | c06 | c05,c06,c07,c08,c13,c14,c15,c16 | 0.4 | 0.444444444 |
| alltp | pgmA | region | pgmA | GS | GS_expanded | 20 | 1 | 12 | 1 | c06 | c05,c06,c07,c08,c13,c14,c15,c16, c21,c22,c23,c24 | 0.6 | 0.6 |
| alltp | ptnABCD | region | ptnABCD | GS | GS_primary | 20 | 2 | 8 | 2 | c06,c08 | c05,c06,c07,c08,c13,c14,c15,c16 | 0.147368421 | 0.275689223 |
| alltp | ptnABCD | region | ptnABCD | GS | GS_expanded | 20 | 2 | 12 | 2 | c06,c08 | c05,c06,c07,c08,c13,c14,c15,c16, c21,c22,c23,c24 | 0.347368421 | 0.434210526 |
